# Impact of variants and vaccination on nasal immunity across three waves of SARS-CoV-2

**DOI:** 10.1101/2024.05.29.596308

**Authors:** Jaclyn M. Long, Vincent N. Miao, Anna H. Owings, Ying Tang, Joshua D. Bromley, Samuel W. Kazer, Kyle Kimler, Chelsea Asare, Carly G. K. Ziegler, Samira Ibrahim, Tasneem Jivanjee, Micayla George, Andrew W. Navia, Riley S. Drake, Adam Parker, Benjamin C. Billingsley, Paul Dotherow, Spurthi Tarugu, Sai K. Kota, Hannah Laird, T. Grant Wichman, Yesenia T. Davis, Neha S. Dhaliwal, Yilianys Pride, Yanglin Guo, Michal Senitko, Jessie Harvey, John T. Bates, Gill Diamond, Michael R. Garrett, D. Ashley Robinson, I.J. Frame, Jonathan J. Lyons, Tanya O. Robinson, Alex K. Shalek, Bruce H. Horwitz, Sarah C. Glover, Jose Ordovas-Montanes

## Abstract

SARS-CoV-2 infection and COVID-19 disease vary with respect to viral variant and host vaccination status. However, how vaccines, emergent variants, and their intersection shift host responses in the human nasal mucosa remains uncharacterized. We and others have shown during the first SARS-CoV-2 wave that a muted nasal epithelial interferon response at the site of infection underlies severe COVID-19. We sought to further understand how upper airway cell subsets and states associate with COVID-19 phenotypes across viral variants and vaccination. Here, we integrated new single-cell RNA-sequencing (scRNA-seq) data from nasopharyngeal swabs collected from 67 adult participants during the Delta and Omicron waves with data from 45 participants collected during the original (Ancestral) wave in our prior study. By characterizing detailed cellular states during infection, we identified changes in epithelial and immune cells that are both unique and shared across variants and vaccination status. By defining SARS-CoV-2 RNA+ cells for each variant, we found that Delta samples had a marked increase in the abundance of viral RNA+ cells. Despite this dramatic increase in viral RNA+ cells in Delta cases, the nasal cellular compositions of Delta and Omicron exhibit greater similarity, driven partly by myeloid subsets, than the Ancestral landscapes associated with specialized epithelial subsets. We found that vaccination prior to infection was surprisingly associated with nasal macrophage recruitment and activation rather than adaptive immune cell signatures. While patients with severe disease caused by Ancestral or Delta variants had muted interferon responses, Omicron-infected patients had equivalent interferon responses regardless of disease severity. Our study defines the evolution of cellular targets and signatures of disease severity in the upper respiratory tract across SARS-CoV-2 variants, and suggests that intramuscular vaccines shape myeloid responses in the nasal mucosa upon SARS-CoV-2 infection.

## Introduction

SARS-CoV-2 has caused over 700 million infections and almost 7 million deaths worldwide since it emerged in late 2019 as documented by the WHO.^1^ Despite the high population prevalence of previous exposures and vaccinations, SARS-CoV-2 continues to spread between individuals. The nasal mucosa is a primary site of infection, viral replication, and transmission for respiratory viruses.^2–6^ Understanding nasal cellular responses to SARS-CoV-2 - and how they differ across variants, vaccination status, and disease trajectories - is key to developing therapeutic and prophylactic strategies against this virus and others.

The emergence of novel variants of concern, including Delta and Omicron, with the ability to evade adaptive immunity has contributed to the continued circulation of SARS-CoV-2.^2,7^ *In vitro* studies of variant-specific biology have demonstrated unique replication and pathogenicity properties.^8–14^ Additionally, analysis of both infectious viral load and viral RNA levels from nasal swabs has revealed differences in shedding and transmission dynamics.^15,16^ While the Delta variant was characterized, in part, by rapid intracellular replication, enhanced fusogenicity, and higher viral load, Omicron demonstrated slower replication, less severe pathology, and higher transmission rates.^8,10,11,16^ Investigation of differences in immune responses to each SARS-CoV-2 variant have largely focused on either intracellular mechanisms of innate immune evasion or alterations in antigen-specific adaptive immune cells in the blood.^17,18^ Whether the differences in variant properties are reflected by differences in *in vivo* nasal cell tropism, or in the cellular immune response within the nose, remains unclear.

SARS-CoV-2 vaccines have dramatically reduced the burden of severe disease and death from COVID-19. Hybrid immunity, the combination of immune memory acquired from both vaccination and infection, can provide even more durable and potent protection than vaccination alone.^19,20^ Local immune memory subsets generated during the response to infection are thought to play a key role in this heightened protection, as is supported by work from mouse models of respiratory infection.^20–24^ Investigation of mucosal immune memory populations has shown consistently that prior infection with SARS-CoV-2 or hybrid immunity results in greater numbers of nasal tissue-resident CD8+ and CD4+ memory T cells (Trm) with enhanced functionality than peripheral vaccination alone.^19,20,25–27^ While the expansion of nasal Trm following vaccination alone has been described,^28^ comparison of mucosal antibodies in vaccinated or convalescent individuals revealed higher levels of SARS-CoV-2 specific IgA in nasal lavage fluid after previous infection^29^ and limited durability following vaccination alone.^20,30^ A mounting body of evidence suggests that lack of establishment of durable, effective mucosal immune memory populations is a limitation of our current vaccination strategies for SARS-CoV-2.^31^ However, given that vaccination does significantly improve peripheral immune responses, respiratory symptoms, and disease severity, we sought to understand its impact on the cellular response in the upper airway during acute infection.

Disease outcomes from SARS-CoV-2 vary widely, from asymptomatic infections to severe respiratory distress.^32,33^ Numerous studies have used high-dimensional technologies to identify correlates of COVID-19 severity in the blood.^34–37^ While these have led to valuable insights, the blood is not always representative of immune responses occurring along the respiratory tract.^21^ Multiple groups, including our own, have identified an interferon response as a critical determinant of disease severity, in the contexts of autoantibodies,^38^ inborn errors of immunity,^39,40^ and in muted airway epithelial responses.^41,42^ However, these studies are focused on infections during the original wave of SARS-CoV-2. Given the complex context of variants, prior infections, and vaccinations that comprises the current situation, it is critical to update our understanding of how these host factors relate to severity.

During the original wave of COVID-19 (April-November 2020), we used scRNA-seq to describe nasal cell composition during SARS-CoV-2 infection, identify potential cellular targets of SARS-CoV-2, and uncover epithelial transcriptional states associated with disease severity.^41^ Another study of nasopharyngeal and bronchial samples collected March-May 2020 described epithelial differentiation pathways upon SARS-CoV-2 infection and suggested heightened immune-epithelial interactions in severe cases.^43^ Comparisons of responses in the pediatric and adult airway to SARS-CoV-2 using samples collected from March 2020-May 2021 provided evidence of differential activation of antiviral pathways in children.^44,45^ Others have applied this method to study features of anosmia in olfactory epithelial biopsies or to identify transcriptional features associated with post-acute sequelae using samples collected months after acute SARS-CoV-2 infection.^46,47^ A recent human challenge study with the pre-alpha SARS-CoV-2 variant focused on the dynamics of responses in the blood and airway.^48^ This unique experimental paradigm enabled the characterization of nasal epithelial and immune cell states during the course of mild infections in unvaccinated individuals. Together, these studies have begun to shed light on the complex cellular response to SARS-CoV-2 in the human nasal mucosa. However, a detailed comparison of responses across three waves of naturally-acquired infection, encompassing major viral variants, vaccines, and disease severity, has yet to be reported.

Here, we present a detailed analysis of the impact of variant and vaccination status on nasal cellular phenotypes across three distinct waves of COVID-19. We integrated nasal scRNA-seq data from 45 participants in our prior study during the original wave of SARS-CoV-2^41^ with 67 additional samples collected during the Delta and Omicron waves. This enabled the identification of detailed epithelial and immune cell states. Comparison across participant groups revealed that while some cellular changes are consistent across variants, such as a decline in resting ciliated population and emergence of interferon-stimulated clusters, other cell states were unique to particular waves. By analyzing the detailed cell type composition of each participant, we found that Delta- and Omicron-responsive cellular “ecosystems”, defined as the collective of diverse cell types present in the tissue, were similar to each other, while Ancestral cases appeared distinct. We observed a striking increase in abundance of SARS-CoV-2 RNA in Delta cases and defined the makeup of SARS-CoV-2 RNA+ cells within each group. When we compared responses between vaccinated and unvaccinated cases, we found that vaccination is associated with nasal macrophage recruitment and gene signatures of inflammation and antigen presentation. Finally, we identified distinct gene signatures and cell states in each variant that were associated with COVID-19 severity. Together, our data provides a comprehensive view of the diversity of nasal cellular responses to SARS-CoV-2, defines cellular targets and potential severity biomarkers across variants, and suggests that intramuscular vaccination may alter the nasal macrophage response.

## Results

### Expanding the cellular atlas of the human nasal mucosa during SARS-CoV-2 infection

Nasopharyngeal swabs were collected from a total of 112 adult patients at the University of Mississippi Medical Center **(Figure 1A-1B, Table 1)**, including 26 control participants who had no respiratory symptoms and tested negative for SARS-CoV-2 by PCR and 86 participants who had positive SARS-CoV-2 PCR tests. We combined data from our previously published study^41^ (13 controls and 32 “Ancestral” cases; April-November 2020) with samples collected during the Delta (13 controls and 33 cases; July-September 2021) and Omicron (21 cases; January-February 2022) waves of the COVID-19 pandemic. The Ancestral and Omicron participants were slightly older (median age 58.5 and 61 years, respectively), than Control (median age 51.5 years) and Delta (median age 50 years) cases, and racial identity also varied across groups **(Table 1)**. Among pre-existing conditions, we observed higher rates of diabetes and inflammatory bowel disease in Controls and of congestive heart failure in Omicron cases **(Table 1)**. Rates of Remdesivir use also varied across variant groups **(Table 1)**. The Delta and Omicron wave cohorts included participants who had received at least 2 doses of an mRNA vaccine for COVID-19 prior to infection as well as participants who were unvaccinated **(Figure 1A, Table 1)**. Within each variant cohort, as well as within vaccination groups, participants experienced a range of COVID-19 severity quantified by a previously described score of required respiratory support (0-8) **(Figure 1C-1D, Table 1, Supplementary Figure 1A. Supplementary Figure 2A).**^49^ All COVID-19 swabs were collected while participants experienced acute respiratory symptoms. Though the timing was variable between individuals, there was no significant difference in the number of days from positive COVID-19 test to sample collection between participant groups **(Supplementary Figure 1B-1D, Supplementary Figure 2B-2D).** Full clinical metadata of participants can be viewed in **Supplementary Table 1.** Comparison of SARS-CoV-2 specific antibody levels in plasma collected at the same time as the nasopharyngeal swab did not reveal any significant differences in IgG titers between disease groups, with the exception of higher levels of nucleoprotein (NP)-specific antibodies in unvaccinated Delta participants compared to vaccinated Delta participants **(Supplementary Figure 1H-1K, Supplementary Figure 2H-2J)**. Due to the timing of sample collection, we were unable to resolve whether anti-NP antibodies were produced in response to a prior or ongoing infection, and thus do not further subdivide our cohort based on serology-confirmed exposure history to SARS-CoV-2.

**Figure 1:**
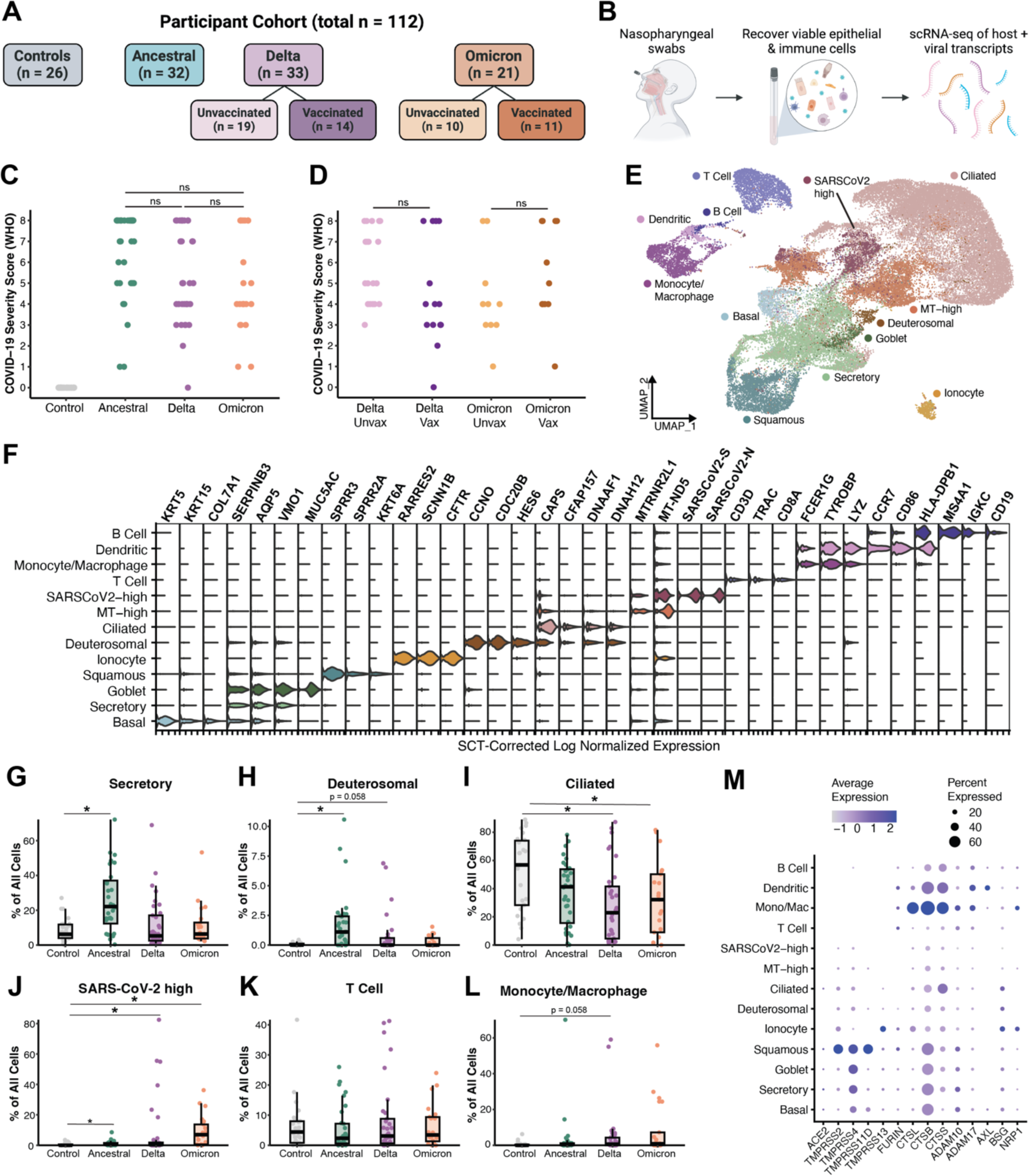
Cellular composition of the nasal mucosa across three SARS-CoV-2 variant waves. (A) Study schematic: participant cohort breakdown (Biorender) (B) Study schematic: nasopharyngeal swabs, cell isolation, and scRNA-seq (Biorender) (C-D) COVID-19 severity, defined by the WHO score of respiratory support required at peak of disease, divided by variant wave (Control, Ancestral COVID-19, Delta COVID-19, and Omicron COVID-19) (C) and variant + vaccination status (D). Statistical test represents Kruskal-Wallis test results across all conditions with Benjamini-Hochberg correction for multiple comparison across all cell types. Statistical significance asterisks represent results from Dunn’s post hoc testing. *p < 0.05 (E) UMAP of 48,730 cells from all participants colored by major cell type following iterative louvain clustering (F) Violin plots of cluster marker genes for cell type annotations shown in (E) (G-L) Frequency of select cell types separated by SARS-CoV-2 variant. Statistical test as in (C-D). *p < 0.05 (M) Expression of SARS-CoV-2 entry factors and proteases across major cell types

**Table 1:**
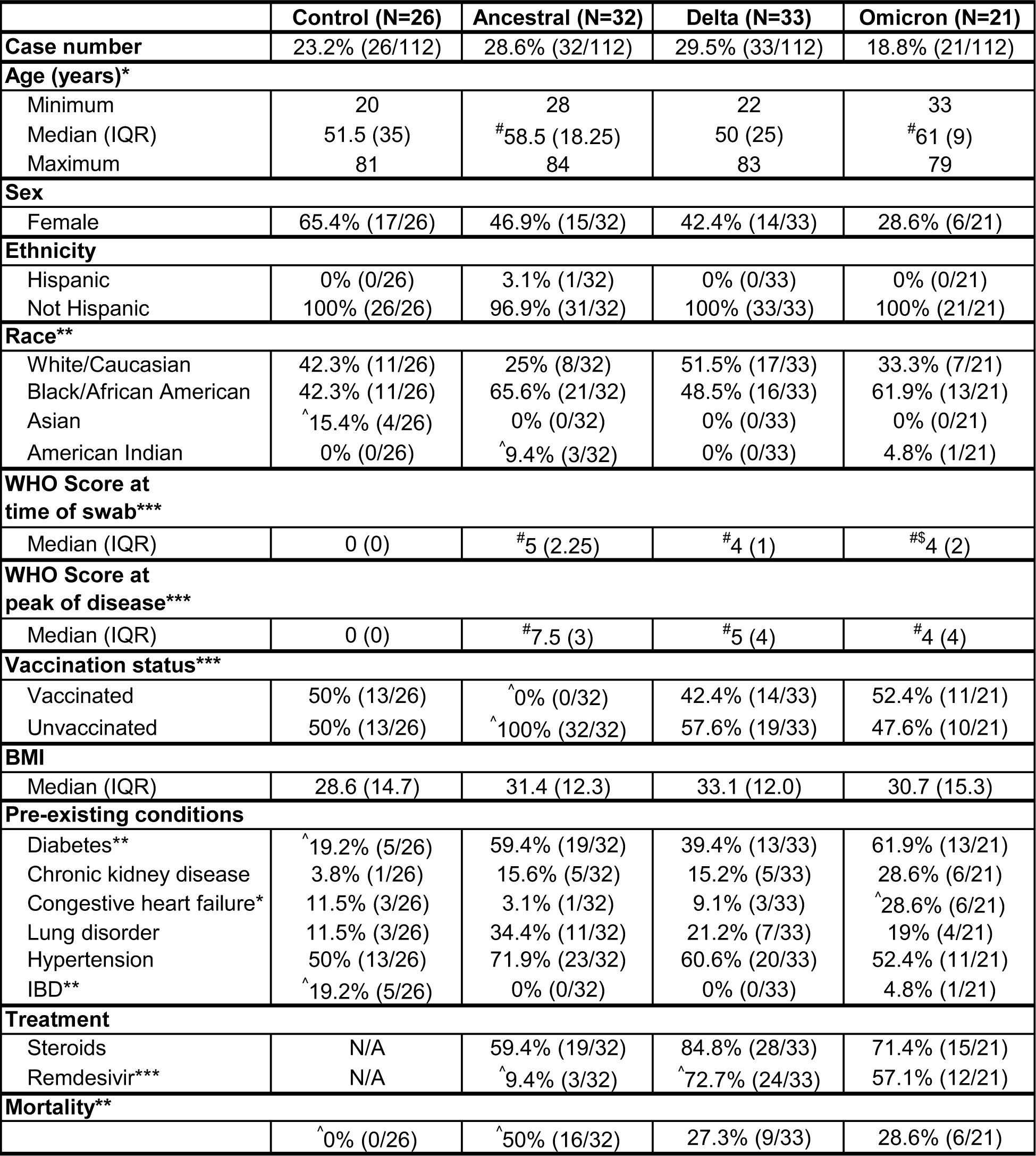
Participant Cohort. Continuous variables were compared by Kruskal-Wallis test. Categorical variables were compared by chi-square test. *p<0.05, **p<0.01, ***p<0.001, otherwise non-significant. ^#^Dunn post-hoc test relative to Control group, p<0.05; ^$^Dunn post-hoc test relative to Ancestral group, p<0.05; ^>^^chi-square standardized Pearson residual >2; otherwise non-significant. IQR: Inter-quartile range; IBD: inflammatory bowel disease.

|  | **Control (N=26)** | **Ancestral (N=32)** | **Delta (N=33)** | **Omicron (N=21)** |
| --- | --- | --- | --- | --- |
| **Case number** | 23.2% (26/112) | 28.6% (32/112) | 29.5% (33/112) | 18.8% (21/112) |
| **Age (years)\*** |  |  |  |  |
| Minimum | 20 | 28 | 22 | 33 |
| Median (IQR) | 51.5 (35) | <sup>#</sup>58.5 (18.25) | 50 (25) | <sup>#</sup>61 (9) |
| Maximum | 81 | 84 | 83 | 79 |
| **Sex** |  |  |  |  |
| Female | 65.4% (17/26) | 46.9% (15/32) | 42.4% (14/33) | 28.6% (6/21) |
| **Ethnicity** |  |  |  |  |
| Hispanic | 0% (0/26) | 3.1% (1/32) | 0% (0/33) | 0% (0/21) |
| Not Hispanic | 100% (26/26) | 96.9% (31/32) | 100% (33/33) | 100% (21/21) |
| **Race\*\*** |  |  |  |  |
| White/Caucasian | 42.3% (11/26) | 25% (8/32) | 51.5% (17/33) | 33.3% (7/21) |
| Black/African American | 42.3% (11/26) | 65.6% (21/32) | 48.5% (16/33) | 61.9% (13/21) |
| Asian | <sup>^</sup>15.4% (4/26) | 0% (0/32) | 0% (0/33) | 0% (0/21) |
| American Indian | 0% (0/26) | <sup>^</sup>9.4% (3/32) | 0% (0/33) | 4.8% (1/21) |
| **WHO Score at time of swab\*\*\*** |  |  |  |  |
| Median (IQR) | 0 (0) | <sup>#</sup>5 (2.25) | <sup>#</sup>4 (1) | <sup>#</sup>4 (2) |
| **WHO Score at peak of disease\*\*\*** |  |  |  |  |
| Median (IQR) | 0 (0) | <sup>#</sup>7.5 (3) | <sup>#</sup>5 (4) | <sup>#</sup>4 (4) |
| **Vaccination status\*\*\*** |  |  |  |  |
| Vaccinated | 50% (13/26) | <sup>^</sup>0% (0/32) | 42.4% (14/33) | 52.4% (11/21) |
| Unvaccinated | 50% (13/26) | <sup>^</sup>100% (32/32) | 57.6% (19/33) | 47.6% (10/21) |
| **BMI** |  |  |  |  |
| Median (IQR) | 28.6 (14.7) | 31.4 (12.3) | 33.1 (12.0) | 30.7 (15.3) |
| **Pre-existing conditions** |  |  |  |  |
| Diabetes** | <sup>^</sup>19.2% (5/26) | 59.4% (19/32) | 39.4% (13/33) | 61.9% (13/21) |
| Chronic kidney disease | 3.8% (1/26) | 15.6% (5/32) | 15.2% (5/33) | 28.6% (6/21) |
| Congestive heart failure* | 11.5% (3/26) | 3.1% (1/32) | 9.1% (3/33) | <sup>^</sup>28.6% (6/21) |
| Lung disorder | 11.5% (3/26) | 34.4% (11/32) | 21.2% (7/33) | 19% (4/21) |
| Hypertension | 50% (13/26) | 71.9% (23/32) | 60.6% (20/33) | 52.4% (11/21) |
| IBD** | <sup>^</sup>19.2% (5/26) | 0% (0/32) | 0% (0/33) | 4.8% (1/21) |
| **Treatment** |  |  |  |  |
| Steroids | N/A | 59.4% (19/32) | 84.8% (28/33) | 71.4% (15/21) |
| Remdesivir*** | N/A | <sup>^</sup>9.4% (3/32) | <sup>^</sup>72.7% (24/33) | 57.1% (12/21) |
| **Mortality\*\*** |  |  |  |  |
|  | <sup>^</sup>0% (0/26) | <sup>^</sup>50% (16/32) | 27.3% (9/33) | 28.6% (6/21) |

Single-cell suspensions of viable epithelial and immune cells were isolated from each nasopharyngeal swab as previously described by our group.^41^ ScRNA-seq libraries were generated using Seq-Well S^3^. After iterative removal of low-quality or low-complexity cells **(Supplementary Figure 3, Supplementary Table 2)**, we generated a dataset of 48,730 quality **(Supplementary Figure 4A-4D, see Methods)** cells encompassing diverse epithelial and immune cell types **(Figure 1E)**. We recovered the majority of cell types described in initial atlases of the airway epithelium (**Figure 1F)**.^50–54^. This included ciliated cells (*CAPS, CFAP157, DNAH12*), which are responsible for mucociliary clearance together with secretory and goblet cells (*AQP5, SERPINB3, VMO1, MUC5AC*), the main producers of mucus in the airway. We identified basal cells (*KRT5, KRT15, COL7A1*), the main stem cells of the airway responsible for regeneration and repair, as well as squamous cells (*SPRR3, SPRR2A, KRT6A)* important in barrier function. Our dataset also includes rare cell types such as recently-described ionocytes (*CFTR, RARRES2, SCNN1B)*, whose function is likely related to ion transport and fluid regulation,^55,56^ and deuterosomal cells (*CCNO, CDC20B, HES6)*, specialized precursors of ciliated cells arising from secretory or goblet cells and characterized by high expression of genes involved in centriole amplification.^57,58^

Within epithelial cells, we recovered clusters of cells with transcriptional patterns such as high amounts of SARS-CoV-2 RNA or mitochondrial reads that drove their transcriptional identity rather than the underlying cell type. These clusters expressed low levels of ciliated genes (e.g., *CAPS*), formed distinct clusters despite the removal of SARS-CoV-2 genes from the principal component analysis, and persisted during iterative rounds of removing low-quality cells **(Supplementary Figure 3)**. Thus, we retained them for further analysis. In addition to epithelial cells, we also recovered multiple immune cell types with important roles in viral infection. The immune compartment included monocytes/macrophages (*FCER1G, TYROBP, LYZ*) dendritic cells (*CCR7, CD86*), T cells (*CD3D, TRAC, CD8A*) and B cells (*MS4A1, IGKC, CD19*).

To understand changes in broad cell types during infection, we compared the frequency of these major cell types across each participant cohort (Control, Ancestral, Delta, or Omicron) **(Figure 1G-1L)**. In Ancestral cases, we observed significantly higher proportions of secretory (Cohen’s d effect size =1.2, p=4.6*10^-4^) and deuterosomal (d=0.92, p=5.2*10^-5^) cells compared to controls, suggesting preferential differentiation towards ciliated cells through the deuterosomal intermediate.^43,57^ In both the Delta (d=0.84, p=7.8*10^-3^) and Omicron (d=0.72, p=0.037) cases, there was a reduction in the average frequency of ciliated cells compared to controls. In all three variants, though to a greater extent in Delta and Omicron, we observed an increase in frequency of SARS-CoV-2 RNA-high cells. Interestingly, with the exception of monocytes/macrophages in the Delta cases, we did not see major changes in the frequency of myeloid or T cells as a fraction of all cells in SARS-CoV-2 cases.

We next evaluated the expression of the SARS-CoV-2 receptor *ACE2* as well as proteases and co-factors important for viral processing **(Figure 1M)**.^59–64^ While *ACE2* was detected at low levels overall, it is highest in secretory, goblet, squamous, and ciliated epithelial cells. *TMPRSS2* is highly expressed in squamous cells, and is also found in other epithelial cells. *TMPRSS4* and *CTSB* are the most highly expressed proteases and are present in multiple cell types. Notably, the SARS-CoV-2 RNA-high cluster had low expression of *ACE2* and many other genes of interest.

### Defining diverse SARS-CoV-2 responsive epithelial and immune cell states in the human nasal mucosa

Next, we interrogated the diversity of cellular states present in the nasal mucosa during the antiviral response. For each major epithelial cell type with at least 1000 cells (basal/secretory, squamous, ciliated, MT-high, and SARS-CoV-2 RNA-high), we separated the cells, re-normalized the data, and performed a new clustering analysis **(Supplementary Figure 5, Supplementary Tables 3-4)**. In the secretory compartment **(Supplementary Figure 5B, 5D)**, multiple clusters were marked by high expression of individual genes, such as lineage markers (*SCGB1A1, MUC5AC*) and keratins (*KRT4, KRT24*). With the exception of *KRT4* as a marker for a basal/secretory transitional subset in the mouse trachea,^56^ these cytokeratins are relatively understudied in the airway compared to others. There was also a notable cluster with high expression of neutrophil recruitment chemokines *CXCL1*, *CXCL2*, and *CXCL8*. Similarly, analysis of the squamous cells **(Supplementary Figure 5C,5D)** revealed a distinct *CXCL8*-high cluster. The ciliated subsets **(Supplementary Figure 5E, 5H)** included two with high expression of cilia-forming genes (*DNAAF1, DNAH12)* and three marked by interferon (IFN) response genes (*IFITM3, IFIT6, ISG15, HLA-A, HLA-B)*. We also observed interesting diversity within the MT-high and SARS-CoV-2 RNA-high clusters **(Supplementary Figure 5F-5H)**. Notably, one SARS-CoV-2 high cluster uniquely expressed circadian regulators *PER1* and *PER2,* as well as the hormone *GDF15* and the transcription factor *EGR1* and *GDF15*. This phenotype is consistent with a set of genes identified as correlated with high amounts of SARS-CoV-2 RNA within nasopharyngeal ciliated cells in a recent SARS-CoV-2 human challenge study.^48^

We applied the same sub-clustering approach to immune cell types with at least 1000 cells and identified multiple subsets of T cells and myeloid cells (**Supplementary Figure 6, Supplementary Table 5).** The strongest signature within T cells was cytotoxicity (*GZMB, GNLY*), with another cluster being marked by high *CXCR4* expression. **(Supplementary Figure 6B, 6D)**. We identified multiple myeloid clusters expressing unique patterns of chemokines: one with *CCL3* and *CCL4*, one with *CXCL5* and *CXCL1*, and one with *CXCL10* and *CXCL11* **(Supplementary Figure 6C-6D)**. There was also one more monocyte-like cluster with high expression of *LYZ*. Across the entire dataset, our final annotation included 57 detailed cell subsets, highlighting the diversity of cellular states in the nasal mucosa during the antiviral response to SARS-CoV-2.

### Alteration of nasopharyngeal cellular composition during distinct waves of SARS-CoV-2 infection

To visualize the relationships between these end clusters, we generated a host cell phylogenetic “tree” using ARBOL **(Figure 2A).**^65^ The centroids of each cellular subset were hierarchically clustered based on human gene expression distances, such that transcriptionally similar subsets would have the shortest distances. As expected, the majority of subsets derived from the same major cell type clustered together. Interesting exceptions included inflammation/NFkB-responsive secretory cells (*NFKBIA+TNFAIP3+*), which clustered closer to squamous cells, and secretory-like (*SERPINB3*+*VMO1+AQP5+*) ciliated cells which were grouped with other secretory clusters, indicating, in these cases, convergent end state transcriptional responses that overshadow original cell type identity programs. In some cell types, such as T cells, the sub-clusters were all closely related to each other, while in others (e.g., ciliated or secretory cells), there was a greater amount of diversity between subsets as reflected by the hierarchical distances. Additionally, participant diversity was maintained in the end clusters in this detailed annotation, indicating relatively little participant-specific effects. This indicates the need to better-understand how a virus such as SARS-CoV-2 can induce unique or intermediate epithelial cell states.

**Figure 2:**
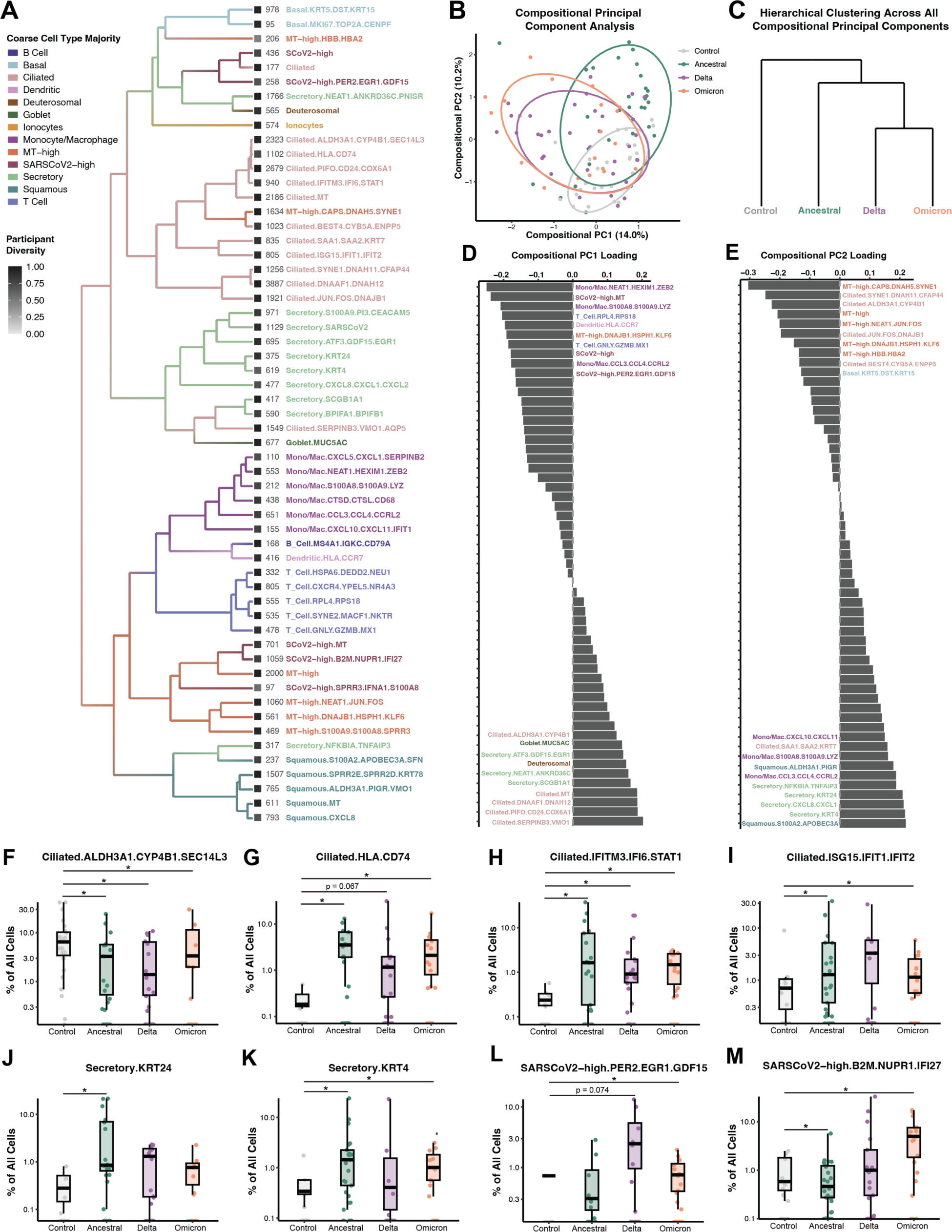
Compositional analysis reveals unique nasal ecosystems in Delta and Omicron relative to Ancestral infections. (A) Cell lineage tree generated using ARBOL depicting relationship between 57 detailed cell subsets. Hierarchical clustering was performed on end clusters identified in Supplementary Figures 2 and 3. Detailed subset labels are colored by major cell type, and branches are colored by the cell type majority for each branch. Squares represent participant diversity (Shannon’s diversity index) for that subset. Cell numbers for each subset are displayed to the left of the label. (B) Compositional principal component analysis of all samples. Each point represents an individual participant and distance reflects differences in abundance of end clusters portrayed in (A). Dots are colored by condition and ellipses for each condition represent centroids. (C) Hierarchical clustering of mean compositional principal component values for each condition (D-E) Detailed subset loadings for PC1 (C) and PC2 (D). Top 10 positive and negative loadings are shown for each PC. Subsets are colored by major cell type (F-M) Frequency of select detailed subsets across conditions. Percentage is shown on a log scale. Statistical test represents Kruskal-Wallis test results across all conditions with Benjamini-Hochberg correction for multiple comparison across all cell types. Statistical significance asterisks represent results from Dunn’s post hoc testing. *p < 0.05

Next, we used compositional analysis to assess the “cellular ecosystem” of each sample. For every participant, we calculated the total cell number-normalized abundance of all 57 detailed clusters. We then performed compositional principal component analysis (PCA) to determine the factors driving the greatest amount of variation in composition across individuals **(Figure 2B)**. We observed that control samples had less variation in nasal cell composition, and that Ancestral cases had a distinct compositional landscape compared to Delta and Omicron, which were highly overlapping. When we investigated the similarity of the conditions by performing hierarchical clustering of centroids for each participant group across all of the principal components **(Figure 2C)**, we found that Delta and Omicron compositions were surprisingly similar to each other.

Since the first two principal components defined the greatest amount of variation, we investigated which detailed cell subsets were contributing the most to each component (**Fig 2D-2E)**. This revealed that the compositions of control participants (positive PC1, negative PC2) were mostly defined by ciliated and MT-high subsets, while the unique compositions of Ancestral cases (positive PC1, positive PC2) were most heavily influenced by particular secretory, squamous, and macrophage subsets. Interestingly, the strongest contributors to the shared ecosystem of Delta and Omicron cases (negative PC1) were immune, including multiple macrophage subsets, dendritic cells, and cytotoxic T cells, as well as SARS-CoV-2 RNA-high subsets. These results suggested to us that the similarity in Delta and Omicron compositions may reflect prior vaccination, higher rates of prior exposure, changes in viral tropism, or combinations of these factors.

We then quantified the frequency of detailed cell subsets across the major participant groups. Control samples had the highest frequency of ciliated cells with a resting/metabolic phenotype (**Figure 2F)**. Across all three variants, there was an increase in frequency in IFN-responsive ciliated subsets **(Figure 2G-2I)**, indicating a conserved shift in ciliated cell phenotypes. The *KRT4*-high and *KRT24-*high secretory subsets contributed positively to compositional PC2 **(Figure 2D)** and were enriched in Ancestral cases (*KRT4*-high: d=0.54, p=3.9*10^-4^; *KRT24*-high: d=0.6, p=9.8*10^-3^) **(Figure 2J-2K).** Delta and Omicron cases shared an increase in frequency of *PER2+EGR1+GDF15*+ SARS-CoV-2 RNA-high cells (Delta: d=0.46, p=0.074; Omicron: d=1.12, p=3.9*10^-4^) **(Figure 2L)**, while only in Omicron cases did an IFN-responsive SARS-CoV-2 RNA-high cluster emerge **(Figure 2M).** Together, we identified patterns in nasal cell composition during SARS-CoV-2 infections across all three waves that suggest a distinct shared ecosystem in Delta and Omicron antiviral responses relative to Ancestral.

### Major SARS-CoV-2 variants have distinct cellular targets and viral RNA abundance patterns

With the emergence of variants and immunological memory, SARS-CoV-2 has been shown to have distinct replication dynamics, symptoms, and transmission rates.^17^ We hypothesized that some of this variation may be reflected in which cell types in the nasopharynx are targeted and the amount of viral RNA that accumulates. We used our previously developed method^41^ to quantify viral transcripts and assign cells as SARS-CoV-2 RNA+. Briefly, we aligned our sequencing data to a joint human and SARS-CoV-2 genome^66^ to quantify viral transcripts in each cell. We then estimated the proportion of ambient RNA contamination for each cell and the amount of SARS-CoV-2 RNA within the non-cell associated wells for each sample. Finally, we used these values to test if the amount of viral RNA within a cell was above the background ambient amount to determine if a given cell was SARS-CoV-2 RNA+ **(Figure 3A)**.

**Figure 3:**
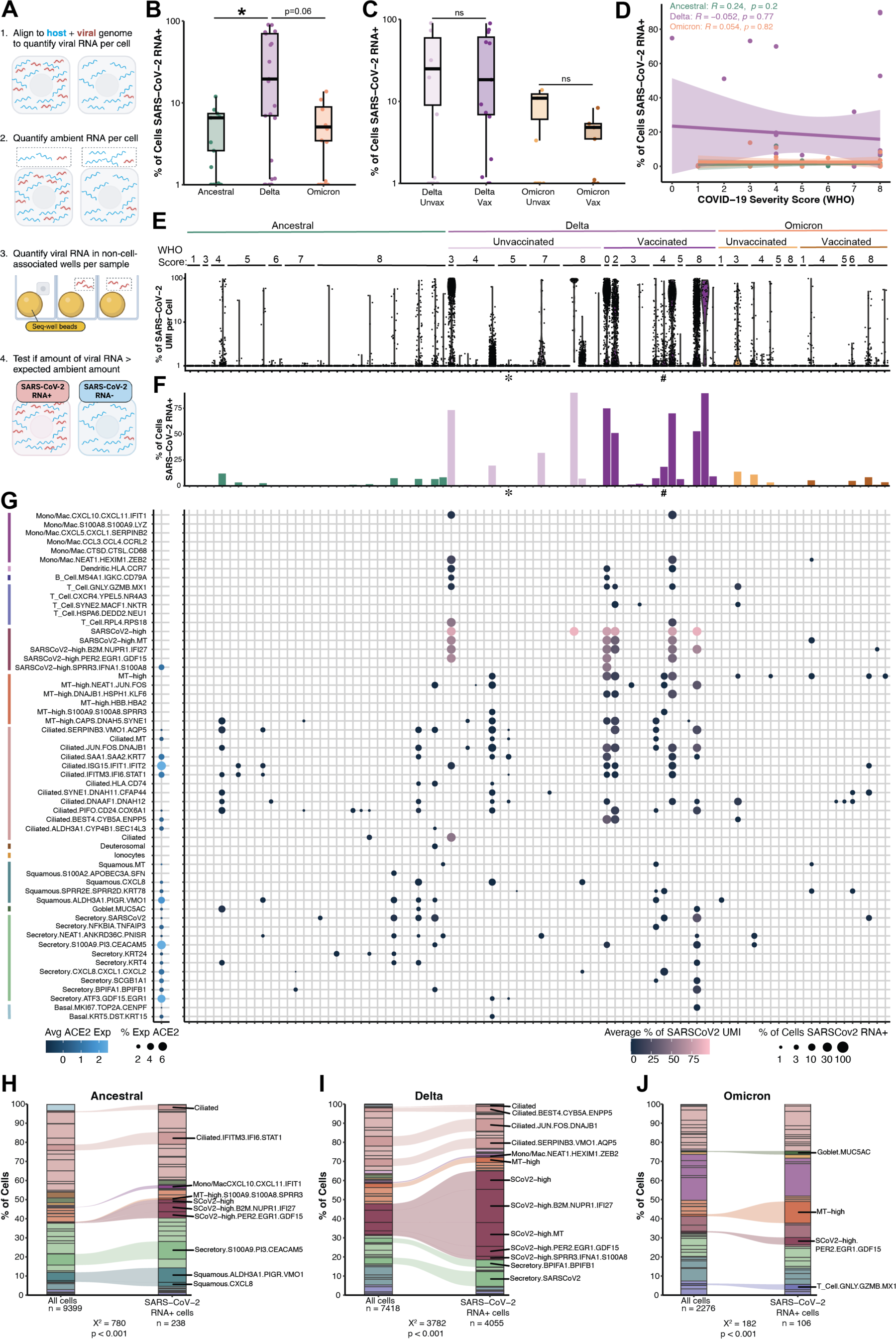
Cellular targets of SARS-CoV-2 in the nasopharynx shift between variants. (A) Schematic of method used to assign SARS-CoV-2 RNA+ cells (B-C) Frequency of SARS-CoV-2 RNA+ cells across variant (B) and vaccination (C) groups. Statistical test represents Kruskal-Wallis test results across all conditions with Benjamini-Hochberg correction for multiple comparison. Statistical significance asterisks represent results from Dunn’s post hoc testing. *p < 0.05 (D) Spearman correlation between COVID-19 severity score (WHO) and frequency of SARS-CoV-2 RNA+ cells across variants. (E) Violin plot of percent of transcripts in each cell that are SARS-CoV-2 transcripts. Each violin represents one participant, ordered by variant, vaccination status, and COVID-19 severity score (WHO). Symbols indicates samples where variants other than Delta were identified from viral sequencing; * = Alpha (B.1.1.7-like); $ = Gamma (P.1-like) (F) Percentage of cells from each participant that were assigned SARS-CoV-2 RNA+ cells. Participants are ordered as in (E). (G) Left: Dotplot of ACE2 expression across detailed cell type (rows). Colored bars represent major cell types. Dot size represents the percentage of cells from that detailed cell type expressing ACE2, and gradient represents average scaled expression. Right: SARS-CoV-2 RNA+ cells for each detailed cell type (rows) and participant (columns, ordered as in (E) an (F)). Dot size represents the percentage of cells from that participant and detailed cell type that are SARS-CoV-2 RNA+. Dots are shown for combinations of participant and detailed cell type with at least 10 cells. Gradient represents the average percent of transcripts in each cell that are SARS-CoV-2 transcripts. (H-J) Distribution of detailed cell types among all cells and SARS-CoV-2 RNA+ cells for Ancestral (H), Delta (I), and Omicron (J) wave samples. Cells for background distribution were restricted to those from participants with SARS-CoV-2 RNA+ cells. In the Delta and Omicron samples, these were further restricted to participants with confirmed Delta (B.1.617.2-like), Omicron (BA.1-like) or Probable Omicron (BA.1-like) variant sequencing results (see Fig S4B-S4C). Chi-squared test was performed on each variant separately. Labeled detailed cell types represent those with chi-squared residual > 2.

We first compared the frequency of SARS-CoV-2 RNA+ cells across our participant groups of interest. We observed a notable increase in SARS-CoV-2 RNA+ cell frequency in Delta cases compared to the other two variants (vs. Ancestral: d=0.79, p=0.018; vs. Omicron: d=0.68, p=0.06) **(Figure 3B)**. This significantly different viral burden was surprising to us given the overlapping cellular compositions of Delta and Omicron cases **(Figure 2B-2C)**. Higher amounts of viral RNA were not necessarily associated with vaccination status nor with peak respiratory support score when divided by variant **(Figure 3C-3D)**.

Next, we evaluated in more detail the total SARS-CoV-2 transcripts and the amount of SARS-CoV-2 RNA+ cells detected for each participant **(Figure 3E-3F).** There was consistent detection of viral transcripts across many participants. For samples collected in the Delta and Omicron waves with sufficient viral RNA material, we performed targeted viral sequencing to determine the precise lineage and sub-lineage **(Supplementary Figure 7A-7B)**. The majority of participants in each wave (Delta: July-September 2021, Omicron: January-February 2022) were confirmed to have the major variant circulating at that time. Two participants sampled in the Delta wave were infected with Alpha (B.1.1.7) or Gamma (P.1.1.2) variants of SARS-CoV-2 **(Figure 3E-3F, * and # symbols, Supplementary Figure 7A-7B)**. While these individuals were included in cellular analyses comparing responses across SARS-CoV-2 waves, they were excluded from direct analysis of cellular targets for each variant **(Figure 3H-3J)**. To ensure that the higher amount of SARS-CoV-2 RNA in Delta wave samples was not an earlier bias in sampling, we compared total viral transcripts across time from symptom onset. Even at matched time points, there was a notable increase in the amount of SARS-CoV-2 RNA in Delta cases **(Supplementary Figure 7C)**. We also investigated whether this could be due to the technical element of total transcripts captured and found that this correlated weakly with total viral transcripts (**Supplementary Figure 7D)**.

To begin to explore the phenotype of SARS-CoV-2 RNA+ cells, we first evaluated *ACE2* expression across all 57 detailed cell subsets **(Figure 3G, left)**. We observed the highest expression of *ACE2* within IFN-responsive ciliated cells (*ISG15*+*IFIT1*+ or *IFITM3*+*STAT1*+), as well as squamous cells with a secretory phenotype (*PIGR*+*VMO1*+), *S100A9*+ secretory cells, and *GDF15+EGR1+* secretory cells. We then evaluated which detailed cell subsets were included in the SARS-CoV-2 RNA+ cells across all 86 cases **(Figure 3G, right)**. Individual participants tended to have SARS-CoV-2 RNA+ cells in multiple distinct subsets, indicating broad cellular targeting. In Ancestral cases, SARS-CoV-2 transcripts were mostly restricted to ciliated, secretory, and squamous cells. Delta cases, on the other hand, also had SARS-CoV-2 RNA transcripts within immune, mitochondrial-high, and SARS-CoV-2-high cells. We detected fewer viral transcripts overall in Omicron samples compared to Delta, and they were mostly restricted to a few subsets, including cilia-high (*DNAAF1*+*DNAH12*+ or *DNAH11*+*CFAP44*+) ciliated cells and *SPRR2E*+*SPRR2D*+ squamous cells. Interestingly, we observed multiple subsets with high amounts of SARS-CoV-2 RNA, despite low or undetectable levels of *ACE2*.

Finally, we asked which cell subsets were significantly enriched in the SARS-CoV-2 RNA+ fraction within each variant by performing a chi-square analysis **(Figure 3H-3J)**. It was necessary to do this on a per-variant basis because of differences in the background distribution of cell types and in the total number of SARS-CoV-2 RNA+ cells detected. The subsets enriched in the viral positive fraction were highly variable across SARS-CoV-2 variants. In Ancestral cases, the enriched cells included IFN-responsive ciliated cells, *S100A9*+ secretory cells, *PIGR+VMO1*+ squamous cells, and select SARS-CoV-2 high clusters. Delta samples had the most dramatic enrichment of SARS-CoV-2 high clusters. Distinct subsets including *BEST4+CYPB5A*+ ciliated cells, secretory-like ciliated cells, and *BPIFA1-*high secretory cells were over-represented in the viral positive cells only in the Delta cases. Omicron cases had the fewest subsets enriched, limited to MT-high cells, goblet cells, *PER1+EGR1*+*GDF15*+ SARS-CoV-2 high cells, and cytotoxic T cells. The only cells that were consistently enriched in viral positive cells across all three variants were the *PER1+EGR1*+*GDF15*+ SARS-CoV-2 high cells, indicating that this may be a conserved transcriptional response to different SARS-CoV-2 variants.

### Vaccination prior to SARS-CoV-2 infection is associated with macrophage activation in the nasal mucosa

In addition to variation across SARS-CoV-2 variants, we sought to assess the impact of prior vaccination on nasal immune responses to SARS-CoV-2. We divided the Delta and Omicron groups into patients who had received at least 2 doses of an mRNA vaccine and those who were unvaccinated. Participant metadata across these groups is shown in **Supplementary Figure 2**. At the time of sample collection, vaccination groups had similar levels of antibodies against the SARS-CoV-2 receptor binding domain (RBD) **(Supplementary Figure 2H-2I)**. During the Delta wave, unvaccinated cases had higher levels of nucleoprotein (NP)-specific antibodies than vaccinated cases, which could reflect an antibody repertoire bias as a result of vaccination or more prior exposures in Delta unvaccinated cases **(Supplementary Figure 2J)**.

First, we compared the frequency of major immune cell types as a percentage of all cells. We found that there were not significant changes in lymphocyte populations, but in Omicron cases, vaccinated participants had higher frequencies of dendritic cells (d=1.36, p=0.073) and monocytes/macrophages (d=1.2, p=0.016) compared to unvaccinated participants **(Figure 4A-4D)**. Intriguingly, though not statistically significant, we also observed trends towards higher frequencies of dendritic cells, monocytes/macrophages, and T cells among vaccinated control cases (collected during the Delta wave) compared to unvaccinated controls (collected during the Ancestral wave) **(Supplementary Figure 8A-8D)**. We did not observe any significant differences in the frequency of major epithelial cell types in vaccinated cases **(Supplementary Figure 8E-8M).** Next, we asked if the influx of macrophages we observed in vaccinated Omicron cases was driven by any individual subsets. We observed increases in frequency of multiple macrophage subsets, including the *CCL3*+*CCL4*+ subset, a cysteine-cathepsin high subset, and a *HEXIM1*+*ZEB1*+ subset, in vaccinated Omicron participants **(Figure 4E-4H)**.

**Figure 4:**
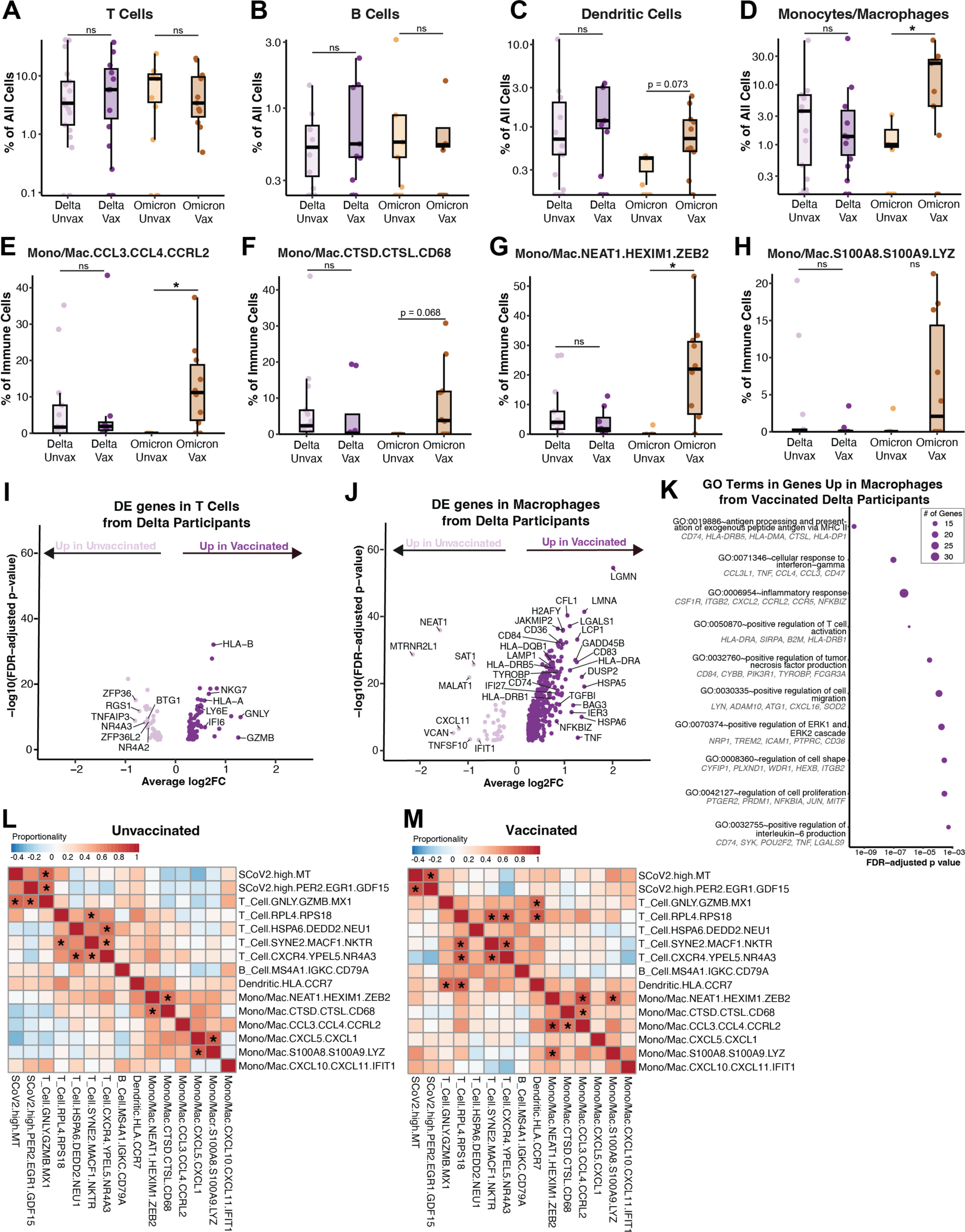
Impact of vaccination on nasal immune responses. (A-D) Frequency of major immune cell types as a percentage of all cells in each sample. Percentage is shown on a log scale. (E-H) Frequency of select detailed immune cell subsets as a percentage of immune cells. Statistical tests for A-H represent Kruskal-Wallis test results across all conditions with Benjamini-Hochberg correction for multiple comparison across all cell types. Statistical significance asterisks represent results from Dunn’s post hoc testing. *p < 0.05 (I-J) Differentially expressed (DE) genes between T cells (I) and macrophages (J) from unvaccinated and vaccinated participants in the Delta cohort. **See Supplementary Table 6** for full list of DE genes. (K) Gene ontology (GO) terms in genes identified as upregulated in macrophages from vaccinated Delta cases in (J). See **Supplementary Table 7** for all identified GO terms. (L-M) Heatmaps of proportionality between abundance of immune cell subsets in unvaccinated (L) and vaccinated (M) cases. *FDR < 0.05

We did not observe similar changes in myeloid cell frequency in Delta cases, but we did have sufficient cells (at least 300 in each group) to perform differential gene expression within immune cell types between cells from vaccinated and unvaccinated participants. This analysis was not done in Omicron cases due to low immune cell numbers in unvaccinated participants. We found that T cells from Delta cases did not have many differentially expressed genes between vaccination groups, but interestingly did upregulate cytotoxicity genes including *GZMB, GNLY*, and *NKG7* in vaccinated participants **(Figure 4I, Supplementary Table 6)**. Surprisingly, we observed a dramatic phenotypic shift in macrophages from vaccinated participants **(Figure 4J, Supplementary Table 6)**. Gene ontology analysis of the upregulated genes revealed pathways including MHC class II antigen presentation, IFNγ response, TNF production, and cell migration **(Figure 4K, Supplementary Table 7)**. Together with the frequency shifts observed in Omicron participants, these data suggest that nasal macrophage recruitment and activation during SARS-CoV-2 infection is associated with prior mRNA vaccination.

We then sought to gain a holistic understanding of how vaccination may shift immune networks in the nasal response to SARS-CoV-2. We performed proportionality analysis to assess relationships between immune cell subsets and ask if any clusters were changing in frequency together **(Figure 4L-4M)**.^67,68^ After evaluating all of the proportional pairs in the dataset that included at least one immune cell cluster, we found that the majority of proportional relationships were between cell clusters within a given major cell type; i.e., T cell subsets are more likely to change with each other than with macrophage subsets. In unvaccinated cases, we also observed proportional relationships between cytotoxic T cells and two SARS-CoV-2 high subsets, including the *PER2*+*EGR1*+*GDF15*+ cluster. In vaccinated cases, dendritic cells varied proportionally with cytotoxic T cells and T cells with a naive/memory phenotype (high in ribosomal genes). These results indicate that while there may be a shift towards more DC-T cell interactions in previously vaccinated individuals, there was not an overall drastic change in nasal immune networks.

### Distinct cell states associated with COVID-19 severity across variants

Lastly, we sought to identify cellular and transcriptional signatures of COVID-19 severity across variants. In prior work from our group and others, an impaired upper respiratory tract IFN response has been associated with severe COVID-19.^41,42^ We therefore began by evaluating expression of IFN stimulated genes (ISGs) across disease severity scores for each variant **(Supplementary Figure 9A-9F, Supplementary Table 8)**. We used previously described gene sets expressed in human nasal basal cells upon treatment with IFNα or IFNγ to evaluate a respiratory epithelial-specific signature.^4,50^ We restricted this analysis to cell types with sufficient representation across multiple disease severity scores in all three variant cohorts, which included ciliated cells, secretory cells, and T cells. We observed similar patterns in Ancestral and Delta cases, where the highest scores were in mild cases and the lowest were among the most severe cases. Interestingly in Delta, we also observed low IFN response scores in participants who were eventually hospitalized but did not receive any oxygen therapy (WHO score 3), followed by a smaller peak in participants who went on to receive non-invasive ventilation or high-flow oxygen (WHO score 5). In Omicron cases, IFN responses were much lower overall, and increased slightly with COVID-19 severity.

To ask whether ISG expression within particular cell types correlated with COVID-19 severity on a participant-by-participant basis, we used pseudobulk analysis. We restricted this comparison to ciliated cells since this cell type had the largest number of participants represented across severity scores for all three variants. We found that in both Ancestral and Delta cases, the expression of the ISGs *STAT1, DDX58, IRF9, CXCL11,* and *NLRC5* in ciliated cells negatively correlated with COVID-19 severity **(Figure 5A-5E)**. In Omicron cases, however, the expression of ISGs in ciliated cells either did not vary with disease severity (*STAT1, DDX58, IRF9, CXCL11)*, or correlated positively (*NLRC5*, *OAS1*) **(Figure 5A-5F)**. Together, these results indicate that in the context of Omicron infections, a robust nasal epithelial IFN response may not be protective against severe COVID-19, and may even be detrimental.

**Figure 5:**
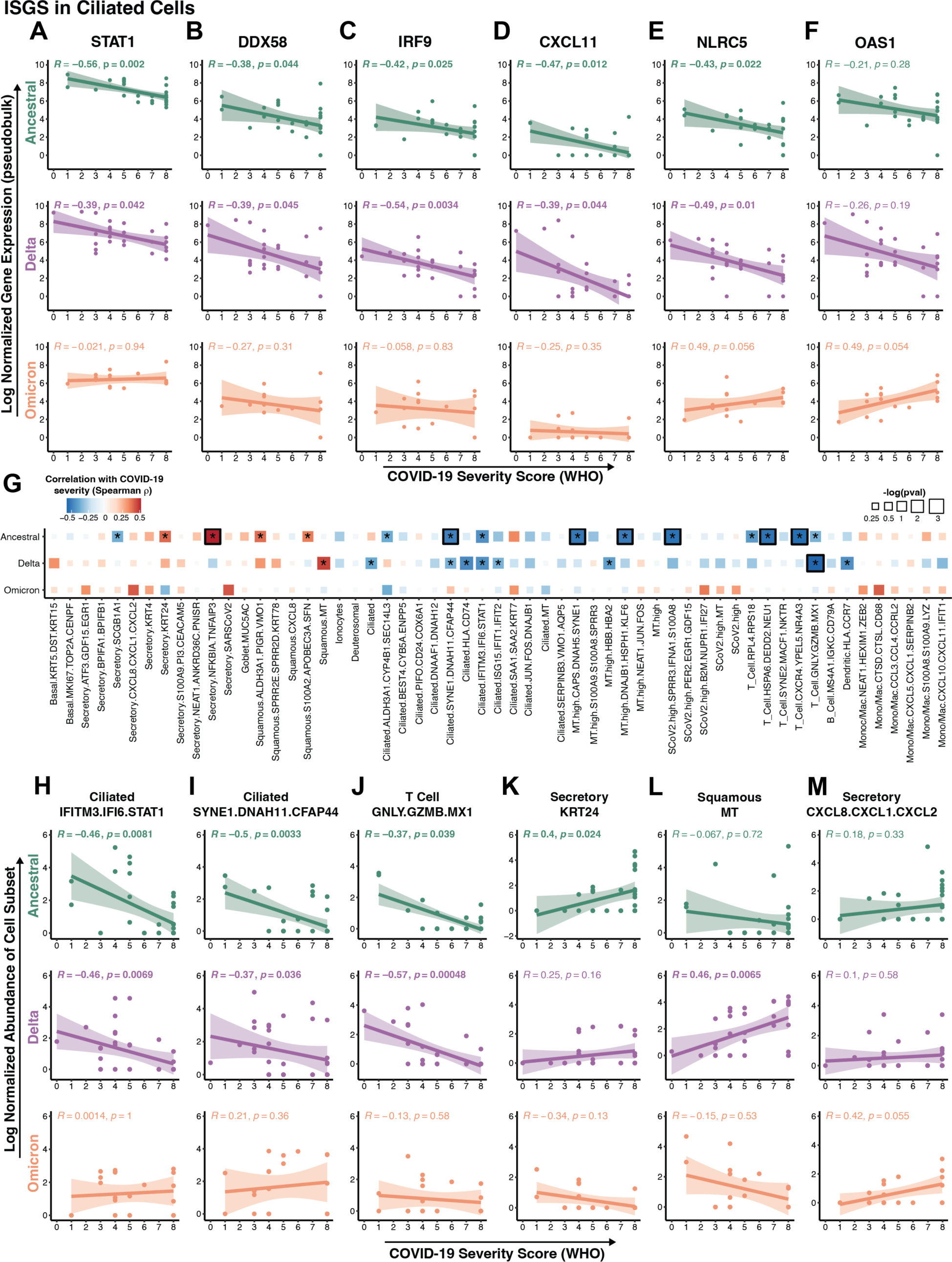
Transcriptional and Cellular Correlates of COVID-19 Severity Across Variants. (A-F) Spearman correlation between COVID-19 severity score and pseudobulk log normalized expression of ISGs (interferon-stimulated genes) in ciliated cells, separated by variant. Each point represents one participant. (G) Heatmap of spearman correlation between COVID-19 severity score and log normalized abundance of select detailed cell subsets, separated by variant. Color indicates spearman *R,* square size indicates -log(pval), border indicates FDR < 0.05, *p<0.05. (H-M) Spearman correlation as in (G) for select detailed cell subsets, separated by variant. Each point represents one participant.

Since the pattern of impaired IFN responses in severe COVID-19 was not consistent across participant groups, we performed a more unbiased analysis of severity correlates. First, we evaluated the relationships between the frequencies of major cell types and SARS-CoV-2 RNA+ cells with all metadata variables, including COVID-19 severity score, within the entire dataset **(Supplementary Figure 10, Supplementary Table 10)**. We found that the frequency of secretory cells and SARS-CoV-2 RNA+ cells was positively correlated with severity, and that the frequency of ciliated cells was negatively correlated, consistent with our observation of more abundant ciliated cells in control samples **(Figure 1I)**. Interestingly, the frequencies of secretory, SARS-CoV-2 RNA+, and squamous cells were also positively correlated with neutrophil levels in the peripheral blood, a known sign of severe lung disease.^32,69^ SARS-CoV-2 RNA+ cells were also correlated with several blood markers, including BUN, sCR, and proBNP.

We observed a positive correlation between vaccination status and dendritic cell frequency, consistent with our observation of nasal myeloid cell activation in vaccinated participants **(Figure 4)**. Because the age of participants varied across groups, we wondered whether any nasal cellular phenotypes might correlate with age. We found a negative correlation between nasal T cell frequency in age, but no differences in any other populations. Investigations of age-related changes in immunity to SARS-CoV-2 infection have similarly identified in T cells in both the periphery and the lung, among other alterations.^70–73^

To determine more precise correlates of severity for each variant, we evaluated whether the abundance of any detailed subsets we identified were correlated with severity score (**Figure 5G-5M)**. There were multiple ciliated cell clusters negatively associated with severity in Ancestral and Delta, but not Omicron, cases, including an IFN-responsive population **(Figure 5G, 5H)** and a *SYNE1*+*DNAH11*+ (cilia-high) population **(5G, 5I).** Cytotoxic T cells were similarly negatively correlated with severity in Ancestral and Delta cases, together with other T cell clusters in Ancestral cases only **(Figure 5G, 5J)**. These results further underscored that components of the immune response correlated with protection in earlier variants may no longer be protective against severe Omicron infections. Other subsets were uniquely correlated with severity in individual variants. In Ancestral cases, two subsets of secretory cells (*KRT24+* and *NFKBIA+ TNFAIP3*+) were positively associated with severity, while one population resembling club cells (*SCGB1A1*+) was associated with mild disease **(Figure 5G, 5K)**. Two squamous populations (*ALDH3A1+PIGR+VMO1+* and *S100A2+APOBEC3A+SFN+*) were also associated with severity in Ancestral cases only, while another MT-high squamous subset was found in more severe Delta cases **(Figure 5G, 5L)**. Omicron cases had fewer subsets that were significantly correlated with severity in either direction. Among the subsets that had the strongest positive correlations were *CXCL8+CXCL1+CXCL2+* secretory cells, SARS-CoV-2 RNA-high secretory cells, and *CTSD+CTSL+CD68+* macrophages. Together, our analysis shows that SARS-CoV-2 variants have distinct nasal cell states associated with disease severity, highlighting the need to further understand this biology as novel variants continue to emerge.

## Discussion

Here, we present a detailed characterization of the human nasal cellular response to SARS-CoV-2 infection across viral variants, vaccination status, and disease severity. We hypothesized that SARS-CoV-2 variants, prior vaccination, and their combination would impact the host response at this site of infection. Further, we sought to understand whether the transcriptional and cellular pathways in the nasal mucosa associated with severe COVID-19 would differ given the additional contexts of variants and vaccination. By defining the diverse cell types, subsets, and states present in the human nasal mucosa during acute SARS-CoV-2 infection, we discovered convergent cellular ecosystems in Delta and Omicron cases that were distinct from those in Ancestral cases. Comparison of responses between vaccinated and unvaccinated individuals revealed that in both Delta and Omicron cases, prior vaccination was associated with nasal macrophage activation during acute infection. In assessing the correlates of severity across variants, we found that an impaired nasal epithelial interferon response is no longer associated with severe COVID-19 in Omicron, and rather that each variant has distinct cellular subsets arising in the most severe cases. Together, our work identifies unique responses in the nasal mucosa–not previously measured in studies of blood or *ex vivo* models–underscoring the importance of studying viral immunity in native human barrier tissues.

The human nasal mucosa is a primary site of entry, replication, and transmission for respiratory viruses. Evidence from our group and others has suggested that robust antiviral responses in the upper respiratory tract are critical in the early restriction of viral infections.^41,42,74^ Further, the nasal mucosa can be a durable niche for resident adaptive immune memory cells.^22^ Despite this, our understanding of the cells that define this tissue and how they form coordinated immune responses remains limited. To inform future vaccines and therapeutics for respiratory viruses, there is an urgent unmet need to develop comprehensive knowledge of mucosal immune responses in the human airway.^21,31^ Recent work towards this end includes a human challenge study with pre-alpha SARS-CoV-2 infection, where the authors describe differences in the mucosal and systemic immune responses between participants with sustained, transient, and abortive infections.^48^ While this study provides critical knowledge, it remains important to understand responses during natural infection. Furthermore, the public health contexts in which SARS-CoV-2 circulates continue to grow more complex, as novel variants and sub-variants emerge, rates of prior vaccinations and prior infections rise, and the type and severity of symptoms shifts. In the face of this complexity, it is critical to understand how these various factors impact antiviral immunity in the nasal mucosa, and to assess these changes at the systems level.

The Delta and Omicron variants of SARS-CoV-2 are known to be distinct phylogenetically, and to vary in properties including replication rate, transmission kinetics, and ability to evade both innate and adaptive immunity.^17,75^ Our assessment of the cellular targets and abundance of viral RNA is consistent with this, as we find a dramatic increase in viral RNA in Delta cases. We also observed distinct cellular subsets enriched in the SARS-CoV-2 RNA+ fraction for each variant, including multiple SARS-CoV-2 RNA-high subsets, *BPIFA1+BPIFB1+* secretory cells, and *VMO1+AQP5+* ciliated cells in Delta cases, and a more restricted group including goblet cells in Omicron cases. Notably, only the *PER2+EGR1+GDF15+* SARS-CoV-2 RNA-high subset was consistently enriched across all three variants. This gene signature is consistent with a set of genes identified as highly correlated with SARS-CoV-2 RNA in the recent SARS-CoV-2 human challenge study.^48^ *PER2*, first described as a regulator of the circadian clock,^76^ has also been shown to modulate expression and activity of TLR9,^77^ while its cooperating protein PER1 can also play an anti-inflammatory role.^78,79^ Circadian oscillations of PER2 expression were observed in club cells from murine lung explants,^80^ and in a model of LPS challenge circadian clock genes in epithelial cells were found to regulate neutrophilic inflammation in the lung.^53,80^ The relationship between circadian rhythms and inflammation in airway epithelial cells during infection will be interesting to pursue further. The metabolic hormone GDF15 has been described as a host tolerance factor during acute inflammation through the induction of hepatic triglyceride metabolism.^81^ GDF15 has also been found to be upregulated along with other nasal cytokines in the nasal mucosa in a long-covid proteomics study.^82^ Interestingly, a recent study suggests that EGR1 is a host restriction factor for SARS-CoV-2 replication.^83^ This conserved transcriptional phenotype of SARS-CoV-2 RNA-high cells may reflect a host cell resilience program, and will be interesting to investigate further.

Despite the differences in viral RNA abundance and distribution, we observe substantial overlap in the nasal cellular compositions between participants with Delta and Omicron infections. This shared ecosystem is driven in part by immune subsets, including *S100A8+S100A9+* and *CCL3+CCL4+* monocytes/macrophages, dendritic cells, T cells with a naive/memory phenotype, and cytotoxic T cells, which could reflect increased immune infiltration due to vaccination and higher rates of prior exposures to SARS-CoV-2. Delta and Omicron cases also shared an increase in SARS-CoV-2 RNA-high subsets, which may reflect shared replicative strategies of the two variants. Meanwhile, the cellular compositions of Ancestral cases were defined by specific types of secretory cells, including *KRT4+* and *KRT24+* subsets, as well as the expansion of deuterosomal cells. *KRT4* has been identified as a marker for suprabasal cells, as well as for basal cells differentiating towards other epithelial states.^43,53,55,56,84,85^ Deuterosomal cells are known to be a precursor population for ciliated cells.^53,57^ These results suggest that the presence of adaptive immune memory may limit regeneration through the deuterosomal path.

A concerted effort has been made in the field to understand how SARS-CoV-2 variants continue to evolve strategies to evade detection and elimination by the innate immune system.^18^ Recent work using *in vitro* models, along with the work of many others, has elucidated precise mechanisms by which SARS-CoV-2 variants interfere with the host response, and importantly highlight the convergent evolution of innate immune suppression.^86–88^ Additional studies of distinct immune responses to Delta and Omicron have focused on cellular and humoral immunity to breakthrough infections in the blood, characterizing the phenotypes and kinetics of antigen-specific adaptive immune cells.^89–91^ A scRNA-seq study of PBMCs from severe Delta cases revealed enrichment of a distinct *IFI30*+ monocyte cluster compared to severe cases infected with Wuhan-like SARS-CoV-2.^92^ These studies provide critical knowledge as to the changes in intracellular antiviral activity and peripheral immune responses to each variant. To our knowledge, our study fills a necessary gap of comparing cellular responses during acute infection at a single-cell level across Ancestral, Delta, and Omicron variants in the respiratory tract.

By comparing nasal immune responses in vaccinated and unvaccinated individuals, we found that nasal macrophages were either increased in frequency or displayed an activated phenotype in those who had received prior vaccination. The majority of previous research on the vaccine-induced immunity to SARS-CoV-2 has been focused on the blood. Important recent work investigating components of vaccine-elicited adaptive immune memory during breakthrough infection revealed that pre-existing antibodies and CD4+ spike-specific memory T cells were the primary early effectors in the recall response.^30^ Given the evidence that infection-induced mucosal immunity can provide stronger protection than peripheral immunity induced by vaccination alone,^19–27^ we expected to see a subtle phenotype in the nasal adaptive immune populations in previously vaccinated individuals. Surprisingly, we found that while the frequencies and gene expression profiles of adaptive immune cells were similar between unvaccinated and vaccinated participants, there was evidence of recruitment and activation of macrophages upon breakthrough infection in vaccinated individuals.

This nasal macrophage phenotype could arise from an increase in Fc receptor mediated activation from vaccine-elicited antibodies or response to cytokines from vaccine-induced memory T cells, but more data is necessary to make these connections.^93^ Additionally, multiple groups have investigated the possibility of trained immunity in myeloid subsets after vaccination.^94–96^ A systems vaccinology study demonstrated increased frequency of inflammatory monocytes in the blood following booster vaccination which correlated with the neutralizing antibody response.^97^ Additional studies have described increased IL-1 production resulting from lipid nanoparticles in vaccine formulations and short-term epigenetic alterations in monocytes following vaccination.^98,99^ Furthermore, naturally-acquired infection with SARS-CoV-2 can impact monocyte training measured in PBMCs for up to one year through enhanced IL-6 activity.^100^ Our data raise the intriguing possibility that the trained immunity–arising from a combination of vaccination and/or prior infection–in peripheral monocytes could result in enhanced macrophage recruitment and activation in the respiratory tract during acute infection. Of note, there was a trend in our cohort towards higher levels of monocytes/macrophages within vaccinated control individuals compared to unvaccinated controls. Further work will be required to investigate this overall model.

By including patients with a range of COVID-19 severity in all of our variant and vaccination groups, we were able to investigate whether nasal cellular and transcriptional correlates of disease severity were consistent across these contexts. Our prior work, together with many others, has shown that in the initial waves of SARS-CoV-2, interruption or impairment of the interferon response was associated with severe COVID-19.^38–42,101^ We found that while the expression of ISGs and abundance of cell subsets including IFN-stimulated ciliated cells was associated with mild symptoms in Ancestral and Delta cases, this was no longer the case with Omicron. Instead, the expression of particular ISGs such as *OAS1* in ciliated cells was positively correlated with COVID-19 severity in Omicron cases, as was the abundance of a *CXCL8+CXCL1+CXCL2+* secretory epithelial subset. These results highlight the need to continue to build knowledge of how immune and inflammatory responses to SARS-CoV-2 relate to disease outcome as novel variants and sub-variants emerge, and individuals are frequently re-infected and boosted to a rapidly-evolving viral species. It will be of interest to understand which specific features of adaptive immunity or inflammatory memory may help to mask muted intrinsic interferon responses.^102^

Much previous and ongoing work has sought to characterize the immune responses of mild and severe patients in a systematic manner. A recent blood-based multi-dimensional study of individuals infected between May and December of 2020 defined cellular and molecular signatures associated with five distinct illness trajectory groups.^37^ Interestingly, this study found that the most severe cases had decreased numbers of natural killer cells and decreased expression of cytotoxicity genes, consistent with our finding that cytotoxic T cells negatively correlated with severity in Ancestral and Delta cases. Additional studies from earlier in the pandemic described aberrant features of the peripheral immune response in patients with severe disease, including inflammatory mediators, distinct monocyte subsets, and mistargeted cytokines.^34–36^ There has yet to be a study comparing these features across SARS-CoV-2 variants, in the blood or the airway.

Together, our study provides a comprehensive view of how the ecosystem of cells in the nasal mucosa responding to SARS-CoV-2 infection changes across viral variants, vaccination status, and COVID-19 severity. This unique dataset sheds light on the complexity and variability of the nasal mucosa antiviral response, and provides critical knowledge that may impact the development of future vaccines and therapeutics for SARS-CoV-2 and other respiratory viruses.

### Limitations of the study

Our study has several limitations. First, the inability to cleanly resolve prior exposure by serology prevents us from distinguishing between vaccine-induced, infection-induced, and hybrid immune memory. Second, as we have sampled only one time point per individual, we are unable to comment on immune response dynamics. A recent human challenge study nicely describes the cell types and transcriptional states in the airway and blood with precise temporal information in known naive individuals.^48^ Third, due to the challenging nature of the samples and difficulties with the experimental work, our data contains lower numbers of, and lower quality cells, than would be expected from more routine samples such as PBMCs. Fourth, we recognize that analysis based on single-cell cluster identification is dependent on upstream processing steps and in part operator bias, which may impact reproducibility and robustness. Finally, we acknowledge that detection of SARS-CoV-2 RNA within a cell may not accurately reflect productive infection, and we were not able to perform additional assays to confirm the location and identity of virally infected cells.

## Acknowledgements

We thank the study participants and their families for enabling this research, the clinical support staff at UMMC for assistance in sample collection, and members of the Shalek, Ordovas-Montanes, Horwitz, and Glover labs for thoughtful discussion and feedback. We sincerely think Arlene Sharpe and all members of the Sharpe laboratory for insightful discussion and feedback as well. We thank Joshua Gould, Katherine Siddle, Bo Li, Stephen Fleming, and the Broad Institute viral-ngs and Cumulus teams for assistance with computational pipelines. This project was made possible in part by grant number 2020-216949 from the Chan Zuckerberg Initiative DAF, an advised fund of Silicon Valley Community Foundation to A.K.S. and J.O.-M. The work performed through the UMMC Molecular and Genomics Facility is supported, in part, by funds from the NIGMS, including the Molecular Center of Health and Disease (P20GM144041-M.R.G), Mississippi INBRE (P20GM103476-A.F.) and Obesity, Cardiorenal and Metabolic Diseases-COBRE (P30GM149404-J.E.H.). J.O.M is a New York Stem Cell Foundation – Robertson Investigator. J.O.M was supported by the AbbVie-Harvard Medical School Alliance, the Richard and Susan Smith Family Foundation, the AGA Research Foundation’s AGA-Takeda Pharmaceuticals Research Scholar Award in IBD – AGA2020-13-01, the HDDC Pilot and Feasibility P30 DK034854, the Leona M. and Harry B. Helmsley Charitable Trust, The Pew Charitable Trusts Biomedical Scholars, The Broad Next Generation Award, The Chan Zuckerberg Initiative Pediatric Networks, The Mathers Foundation, The New York Stem Cell Foundation, NIH R01 HL162642, NIH R01 DE031928 and The Cell Discovery Network, a collaborative funded by The Manton Foundation and The Warren Alpert Foundation at Boston Children’s Hospital. J.M.L. was supported by NIH Training Grant 5TL1TR002543; S.W.K. by The Cancer Research Institute’s Irvington Postdoctoral Fellowship; B.H.H. and S.C.G. by NIH R01 DE031928; and A.K.S. by the Bill and Melinda Gates Foundation, a Sloan Fellowship in Chemistry, the NIH (5U24AI118672), and the Ragon Institute of MGH, MIT and Harvard.

## Author Contributions

Conceptualization, J.O.-M., A.K.S., S.C.G., B.H.H.;

Methodology, J.M.L., V.N.M., A.H.O., Y.T., J.D.B., S.W.K., K.K., C.A., C.G.K.Z., A.W.N., S.C.G., J.O.-M., A.K.S., B.H.H.;

Software, K.K., S.W.K.;

Formal Analysis, J.M.L., V.N.M., A.H.O., Y.T., J.D.B., S.W.K., K.K;

Investigation, J.M.L., V.N.M., A.H.O., Y.T., J.D.B., C.G.K.Z., A.W.N., S.I., T.J., M.G., R.S.D., J.T.B., B.C.B., D.A.R.;

Resources, S.C.G., A.H.O.;

Data curation, A.H.O, Y.T., N.S.D., T.O.R., J.M.L.;

Writing-original draft, J.M.L., J.O.-M., A.K.S., S.C.G., B.H.H.;

Writing-review & editing, J.M.L., V.N.M., A.H.O., Y.T., J.D.B., S.W.K., K.K., C.A., C.G.K.Z., S.I., T.J., M.G., A.W.N., R.S.D., A.P., B.C.B., P.D., S.T., S.K.K., H.L., T.G.W., Y.T.D., N.S.D., Y.P., Y.G., M.S., J.H., J.T.B., G.D., M.R.G., D.A.R., I.J.F., J.J.L., T.O.R., J.O.-M., A.K.S., S.C.G., B.H.H.;

Visualization, J.M.L., Y.T.;

Supervision, J.O.-M., A.K.S., S.C.G., B.H.H., T.O.R.;

Project Administration, Y.P., T.O.R.;

Funding Acquisition, J.O.-M., A.K.S., S.C.G., B.H.H.

## Declaration of Interests

V.N.M. reports compensation from MPM Capital and RA Capital Management unrelated to this work. S.W.K. reports compensation for consulting services with Monopteros Therapeutics, Flagship Pioneering, and Radera Biosciences. M.S. reports compensation for consulting services with GE Precision Healthcare LLC. J.J.L. reports compensation for consulting services with Blueprint Medicines and Human Immunology Biosciences, Inc. J.O.M. reports compensation for consulting services with Cellarity, Tessel Biosciences, and Radera Biotherapeutics. A.K.S. reports compensation for consulting and/or SAB membership from Merck, Honeycomb Biotechnologies, Cellarity, Hovione, Ochre Bio, Third Rock Ventures, Relation Therapeutics, Dahlia Biosciences, Bio-Rad Laboratories, IntrECate Biotherapeutics, FogPharma, and Passkey Therapeutics. C.G.K.Z., V.N.M., A.H.O., A.W.N., Y.T., J.D.B., A.K.S., S.C.G., B.H.H., and J.O.-M. are co-inventors on a provisional patent application relating to methods of stratifying and treating viral infections.

## Inclusion & Ethics

We worked to ensure gender balance in the recruitment of human subjects. We worked to ensure ethnic or other types of diversity in the recruitment of human subjects. While citing references scientifically relevant for this work, we also actively worked to promote gender balance in our reference list. The author list of this paper includes contributors from the location where the research was conducted who participated in the data collection, design, analysis, and/or interpretation of the work.

## Methods

### Participant Details

Eligible participants were recruited from outpatient clinics, medical surgical units, intensive care units (ICU), or endoscopy units at the University of Mississippi Medical Center (UMMC) between April 2020 and September 2022. The UMMC Institutional Review Board approved the study under IRB#2020-0065. All participants or their legally authorized representative provided written informed consent. Participants were eligible for inclusion in the COVID-19 group if they were at least 18 years old, had a positive nasopharyngeal swab for SARS-CoV-2 by PCR, had COVID-19 related symptoms including fever, chills, cough, shortness of breath, and sore throat, and weighed more than 110 lb. Participants were eligible for the Control group if they were at least 18 years old, had a current negative SARS-CoV-2 test (PCR or rapid antigen test), and weighed more than 110 lb. Exclusion criteria for the cohort included a history of blood transfusion within 4 weeks and subjects who could not be assigned a definitive COVID-19 diagnosis from nucleic acid testing. 112 individuals were included in the study: 32 individuals sampled during the Ancestral wave of COVID-19 (April-November 2020; 15 female, 17 male), 33 individuals during the Delta wave (July-September 2021; 14 female, 19 male), 21 during the Omicron wave (January-February 2022; 6 female, 15 male) and 26 Control participants (17 female, 10 male). The median age of Ancestral strain participants was 58.5, of Delta participants was 50, of Omicron was 61 and of Control participants was 53 years old. Participants were assigned to the vaccinated group if they had received at least two doses of an mRNA vaccine for COVID-19 prior to sample collection, and to the unvaccinated group if they had not received any COVID-19 vaccine. COVID-19 participants were classified according to the 8-level ordinal scale proposed by the WHO representing the amount of respiratory support required at the peak of their disease.^49^ Full clinical metadata is provided in **Supplementary Table 1**.

### Sample Collection

Nasopharyngeal samples were collected by a trained healthcare provider using FLOQSwabs (Copan flocked swabs) following the manufacturer’s instructions. Collectors would don personal protective equipment (PPE). The nasopharyngeal (NP) swabs were collected from patients in the standard clinical fashion. The swab was then placed into a cryogenic vial with 900 uL of heat inactivated fetal bovine serum (FBS) and 100 uL of dimethyl sulfoxide (DMSO). Vials were placed into a Mr. Frosty Freezing Container (Thermo Fisher Scientific) for optimal cell preservation. A Mr. Frosty containing the vials was placed in a cooler with dry ice for transportation from patient areas to the laboratory for processing. Once in the laboratory, the Mr. Frosty was placed into an 80°C freezer overnight, and on the next day, the vials were moved to liquid nitrogen storage containers.

### Detection of SARS-CoV-2 specific antibodies

Plasma from a subset of participants (14 Control, 25 Ancestral, 29 Delta, 17 Omicron) was collected on the same day as nasopharyngeal swab. SARS-CoV-2-specific antibodies were detected using an ELISA with recombinant SARS-CoV-2 proteins. Briefly, 384-well Maxi-Sorp ELISA plates (ThermoFisher Nunc) were coated with recombinantly expressed protein at a concentration of 3 μg per ml and incubated overnight at 4 °C. Plates were washed 4x with a BioTek 405TS plate washer and blocked with 1% powdered milk solution in PBS for 1 hour at room temperature. Samples were serially diluted in 1% powdered milk/PBS/Tween and applied to the plate. Plates were incubated for 2 hours at room temperature before washing 4x. Horse radish peroxidase-conjugated anti-human IgG or IgM (Southern Biotech) was applied to the plates and incubated for 1 hour at room temperature. Plates were washed 6x, and developed with tetramethylbenzidine (Sigma) for 30 minutes. The reaction was stopped by the addition of 2N H2SO4 and absorbance was read at 450 nm. The endpoint titer was calculated as the inverse of the dilution of sample that produced an absorbance of 0.2 absorbance units above background. All assays were performed in triplicate.

### Dissociation and Collection of Viable Single Cells from Nasopharyngeal Swabs

Swabs in freezing media (90% FBS/10% DMSO) were stored in liquid nitrogen until immediately prior to dissociation as previously described. A detailed sample protocol can be found here: https://protocols.io/view/human-nasopharyngeal-swab-processing-for-viable-si-bjhkkj4w.html.^103^ This approach ensures that all cells and cellular material from the nasal swab (whether directly attached to the nasal swab, or released during the washing and digestion process), are exposed first to DTT for 15 min, followed by an Accutase digestion for 30 min. Briefly, nasal swabs in freezing media were thawed, and each swab was rinsed in RPMI before incubation in 1 mL RPMI/10 mM DTT (Sigma) for 15 min at 37°C with agitation. Next, the nasal swab was incubated in 1 mL Accutase (Innovative Cell Technologies) for 30 min at 37°C with agitation. The 1 mL RPMI/10 mM DTT from the nasal swab incubation was centrifuged at 400 g for 5 min at 4°C to pellet cells, the supernatant was discarded, and the cell pellet was resuspended in 1 mL Accutase and incubated for 30 min at 37°C with agitation. The original cryovial containing the freezing media and the original swab washings were combined and centrifuged at 400 g for 5 min at 4°C. The cell pellet was then resuspended in RPMI/10 mM DTT and incubated for 15 min at 37°C with agitation, centrifuged as above, the supernatant was aspirated, and the cell pellet was resuspended in 1 mL Accutase, and incubated for 30 min at 37°C with agitation. All cells were combined following Accutase digestion and filtered using a 70 um nylon strainer. The filter and swab were washed with RPMI/10% FBS/4 mM EDTA, and all washings combined. Dissociated, filtered cells were centrifuged at 400 g for 10 min at 4°C, and resuspended in 200 uL RPMI/10% FBS for counting. Cells were diluted to 20,000 cells in 200 uL for scRNA-seq. For the majority of swabs, fewer than 20,000 cells total were recovered. In these instances, all cells were input into scRNA-seq.

### Viral Variant Sequencing

RNA was isolated from supernatants collected during swab dissociation using the QIAamp MinElute Virus Spin Kit (QIAGEN Cat. No. 57704) for RNA extraction. The input volume was 100uL of supernatant, and the final elution volume was 50uL. 10uL of each sample was used for PCR detection of SARS-CoV-2 genes. Samples with a CT value < 28 were sent to the University of Mississippi Medical Center Molecular and Genomics Core Facility and sequencing was performed using the Illumina COVIDseq Test and platform. COVIDseq is an amplicon-based next generation sequencing (NGS) test based on 2019-nCoV primers designed to detect RNA from the SARS-CoV-2 virus (based on ARTIC multiplex PCR protocol v4, with 98 amplicons), along with internal control consisting of 11 human mRNA targets. Individual samples were processed and barcoded through the COVIDseq library prep kit (per manufacturer instruction) using an automated Perkin Elmer Zephyr G3 NGS instrument. Subsequently, samples were pooled (96-384 samples) and run on NextSeq500 using MO flow-cell (150 cycle) or HO flow-cell (150 cycle) depending on the number of samples in the pooled library. The DRAGEN COVIDseq Test App (via Illumina BaseSpace Cloud Computing Platform) was used for alignment of reads to confirm positive samples as well identify variants/clade.

### Single Cell RNA-Sequencing

Seq-Well S3 was run as previously described.^41,50,104–107^ Briefly, a maximum of 20,000 single cells were deposited onto Seq-Well arrays preloaded with a single barcoded mRNA capture bead (ChemGenes) per well^108^. Cells were allowed to settle by gravity into wells for 10 min, after which the arrays were washed with PBS and RPMI, sealed with a semi-permeable membrane for 30 min, and incubated in lysis buffer (5 M guanidinium thiocyanate/1 mM EDTA/1% BME/0.5% sarkosyl) for 20 min. Arrays were then incubated in a hybridization buffer (2M NaCl/8% v/v PEG8000) for 40 min, and then the beads were removed from the arrays and collected in 1.5 mL tubes in wash buffer (2M NaCl/ 3 mM MgCl2/20 mM Tris-HCl/8% v/v PEG8000). Beads were resuspended in a reverse transcription master mix, and reverse transcription, exonuclease digestion, second-strand synthesis, and whole transcriptome amplification were carried out as previously described. Libraries were generated using Illumina Nextera XT Library Prep Kits and sequenced using NovaSeq S4 kits at the Broad Institute Sequencing Core: read 1: 21 (cell barcode, UMI), read 2: 50 (digital gene expression), index 1: 8 (N700 barcode).

### Data Preprocessing

Libraries were aligned using STAR within the Drop-Seq Computational Protocol (https://github.com/broadinstitute/ Drop-seq) and implemented on Cumulus (https://cumulus.readthedocs.io/en/latest/drop_seq.html, snapshot 11, default parameters) ^108^. A previously developed custom reference of the combined human GRCh38 (from CellRanger version 3.0.0, Ensembl 93) and SARS-CoV-2 RNA genomes was used for alignment^41^. Aligned cell-by-gene matrices for each array were merged into one Seurat object across all participants. Following CellBender correction, cells were filtered to remove those with fewer than 200 UMI, fewer than 150 unique genes, and greater than 50% mitochondrial reads. This resulted in a dataset of 32994 genes and 55319 cells across 112 study participants (26 SARS-CoV-2-, 86 SARS-CoV-2+)

### SARS-CoV-2 RNA+ Assignment

Cells were assigned as positive for SARS-CoV-2 RNA using the procedure described in Ziegler et al. 2021. We first aligned to a custom reference genome containing both human and SARS-CoV-2 genes and quantified the total number of SARS-CoV-2 UMI in each cell. Next, we determined the proportion of ambient RNA in each cell by comparing the raw matrices with CellBender-corrected matrices. CellBender (v0.2.0) remove-background function was run with default parameters and epochs = 50, expected_cells = 500, hardware_boot_disk_size_GB = 100, hardware_disk_size_GB = 100, low_count_threshold = 5, and total_droplets_included = 10000. Including the top 10,000 barcodes enabled quantification of viral reads in empty wells, since at least 70% of the cell barcodes recovered were low complexity. Finally, we tested whether the abundance of viral RNA in each cell was significantly above the abundance in empty wells given the ambient RNA profile of each array using an exact binomial test (binom.test in R). We performed a Benjamini-Hochberg correction to determine false discovery rate(p.adjust, method = “BH” in R), and cells with FDR < 0.01 were assigned SARS-CoV-2 RNA+.

### Cell Clustering and Annotation

After SARS-CoV-2 RNA+ assignment, CellBender-corrected matrices were used for further analysis. All analysis was carried out using the Seurat (v4.3.0) package in R (v4.2.3). Data was normalized using SCTransform (v0.3.5).^109^ Principal component analysis was run using only human genes. Harmony (v0.1.1) integration was used to account for variant-specific batch effects.^110^ RunHarmony() function was run with default parameters, group.by.vars = “Cohort” and assay.use = “SCT”, where “Cohort” was either Ancestral, Delta, or Omicron. The output dimensionality reduction from Harmony was then used to generate the nearest neighbor graph and Uniform Manifold Approximation and Projection (UMAP). Cells were clustered using Louvain clustering. Unique marker genes for each cluster were identified using Seurat’s FindAllMarkers() function with cutoffs min.pct = 0.25 and logfc.threshold = 0.25.

Prior to annotation, cells were iteratively clustered to remove low-quality, low-complexity cells as well as doublets **(Supplementary Figure 3, Supplementary Table 2)**. Low-quality and low-complexity cells were identified by high expression of mitochondrial genes and little to no expression of other cell type marker genes. Doublets were identified by expression of marker genes for multiple distinct cell types and high feature counts. The first round of clustering the entire dataset (dims = 1:28, resolution = 0.8) resulted in removal of 4005 cells. Upon a second round of clustering the entire dataset (dims = 1:27, resolution = 0.8), all mitochondrial-high clusters expressed some levels of ciliated cell markers, so these were not removed. Clusters were then assigned a lineage of either “Epithelial” (44,952 cells) or “Immune” (6,362 cells).

Epithelial cells were then isolated, normalized with SCTransform, integrated with Harmony, and re-clustered (dims = 1:30, resolution = 1). Preliminary coarse annotations were then assigned using marker genes from the literature.^41,50,52,53,57^ We identified 9 major epithelial cell types, including basal, secretory, goblet, squamous, ciliated, and deuterosomal cells, ionocytes, an MT-high cluster, and a SARS-CoV-2 RNA-high cluster. Each major epithelial cell type was first isolated and re-clustered to ensure that the coarse annotations were correct, and that no low-quality cells or doublets remained **(Supplementary Table 3)**. This resulted in removal of an additional 1,730 epithelial cells. In the second round of epithelial sub-clustering, final annotations were assigned **(Supplementary Figure 5, Supplementary Table 4)**. Clustering parameters were selected such that stable end clusters were identified, each cluster had a minimum of 95 cells, and each cluster had at least 10 significantly differentially expressed genes (FDR < 0.001). Cell subsets were labeled by their coarse annotation and 2-3 distinctive marker genes (e.g. Ciliated.ISG15.IFIT1.IFIT2 denotes an *ISG15*+*IFIT1*+*IFIT2*+ subset of ciliated cells). In the case that there were not unique genes marking clusters beyond the major cell type marker genes, the coarse annotation was maintained.

Sub-clustering basal and secretory cells together revealed 2 basal subsets, 9 secretory subsets, and 1 goblet cell cluster. Basal cells were divided into those expressing classical basal cell markers (*KRT5, KRT15, COL7A1, DST*) and a proliferating cluster (*MKI67, TOP2A, CENPF*). The goblet cell cluster uniquely expressed high levels of *MUC5AC*. Marker genes of the 9 secretory clusters included keratins (*KRT4*, *KRT24*), club cell-like genes (*SCGB1A1*), antimicrobial secretory genes (*BPIFA1, BPIFB1, S100A9, PI3*), chemokines (*CXCL8, CXCL1, CXCL2*), growth-response genes (*GDF15*, *ATF3*, *EGR1*), NF-kB response genes (*NFKBIA, TNFAIP3*), and SARS-CoV-2 genes. We identified 5 squamous clusters marked by expression of small proline-rich proteins (*SPRR2E*, *SPRR2D*), chemokines (*CXCL8*), secretory genes (*PIGR, VMO1*), mitochondrial genes, or the combination of *S100A2* and *APOBEC3A*. Ciliated cells had the greatest amount of diversity with 13 distinct subsets. 3 ciliated clusters had higher expression of ciliogenesis genes (*PIFO, DNAH11, CFAP44, DNAAF1, DNAH12*). 3 other ciliated clusters expressed interferon-response genes (*ISG15, IFIT1, IFIT2, IFITM3, IFI6, STAT1*) or high amounts of MHC class-I antigen presentation genes (*HLA, CD74*). We also identified ciliated clusters marked by cytochrome genes (*CYP4B1*, *CYPB5A*), alarmins (*SAA1, SAA2*), stress-response genes (*JUN, FOS*), and secretory genes (*VMO1, AQP5*). We observed further diversity within both the mitochondrial-high and SARS-CoV-2 high subsets. Within the MT-high cells, there were clusters defined by high expression of ciliogenesis genes, calprotectin subunits (*S100A8*, *S100A9*), hemoglobin genes (*HBB, HBA2*), and additional stress-response genes (*HSPH1*, *NEAT1*). SARS-CoV-2 high clusters were marked by *IFNA1*, circadian clock and growth response genes *(PER1/*2, *EGR1*, *GDF15*), and interferon-response genes (*B2M, IFI27)*. We did not further sub-cluster deuterosomal cells or ionocytes due to low cell numbers.

We applied the same approach to identify diverse subsets within the 6,362 immune cells in our dataset **(Supplementary Figure 6, Supplementary Table 5)**. Removal of immune-epithelial doublets and a low-quality immune cluster resulted in a final immune object with 5,408 cells. The majority of immune cells were either T cells, monocytes, or macrophages, with small populations of B cells and dendritic cells. We further sub-clustered the T cells, and identified clusters marked by ribosomal genes (indicative of quiescence), stress-response genes (*HSPA6, DEDD2*), cytoskeletal gene *MACF1* and natural-killer cell triggering receptor *NKTR*, chemokine receptor *CXCR5*, and cytotoxicity genes (*GNLY*, *GZMB*). We also observed diversity within the monocyte/macrophage compartment. Genes identifying myeloid clusters included the long non-coding RNA *NEAT1*, cysteine-cathepsin genes (*CTSD*, *CTSL*), chemokines (one cluster with *CCL3, CCL4, CCRL2*, one with *CXCL5, CXCL1*, and one with *CXCL10, CXCL11*), and antimicrobial genes (*S100A8, S100A9, LYZ)*. B cells were not sub-clustered due to low cell numbers.

### Compositional Analyses

We used ARBOL (v4.0.0) to understand relationships between our end clusters and their distribution across participant groups.^65^ To create the binary tree, we ran ARBOLcentroidTaxonomy() with default parameters, tree_reduction = “harmony”, centroid_method = “mean”, distance_method = “cosine”, hclust_method = “complete”, and nboot = 100. ggraph (v2.1.0) was then used to plot the phylogenetic tree, where line and text colors represent the coarse annotation majority and the participant diversity and number of cells is shown for each end cluster **(Figure 2A)**.

We then created a log-normalized abundance matrix for further compositional analysis.^67,111^ Abundance was calculated by deriving the frequency of each cluster in each participant and scaling to 500 cells per participant. Principal component analysis on the log-normalized abundance matrix was done using the prcomp() function in R. For hierarchical clustering, we calculated the distance between the centroid for each condition and used the dendro_data() function to extract and plot the dendrogram.

### Differential Gene Expression by Participant Group

To compare gene expression between cells from distinct participant groups we used a wilcoxon rank sum test. Cells from each cell type belonging to either unvaccinated or vaccinated participants were compared in a pairwise manner, implemented using the Seurat FindAllMarkers function **(Supplementary Table 6)**. We considered genes as differentially expressed with an FDR-adjusted p value < 0.001 and log fold change > 0.25. Gene ontology analysis was run using the Database for Annotation, Visualization, and Integrated Discovery (DAVID) **(Supplementary Table 7)**.^112^

### Proportionality Analysis

Proportionality analysis was performed using the propr package (v4.3.0)^113^ on non-log-transformed immune cluster abundances across Delta and Omicron participants. Proportionality was calculated separately for vaccinated and unvaccinated participants. Investigating differential proportionality using the propd() function revealed no clusters with significantly different relationships between unvaccinated and vaccinated groups.

### IFN Gene Module Scores

Gene lists corresponding to ‘‘IFNα Response” or “IFNγ Response’’ are derived from previously published population RNA-seq data from nasal epithelial basal cells treated in vitro with 0.1 ng/mL – 10 ng/mL of IFNα or IFNγ for 12 h **(Supplementary Table 8)**.^4^ Module scores were calculated using the Seurat function AddModuleScore() with default inputs.

### Pseudobulk Analysis

Pseudobulk count matrices for each combination of participant and major cell type were generated using the Seurat Aggregate Expression() function. Analysis was restricted to cell types where at least 3 distinct COVID-19 severity scores had at least 3 participants with at least 10 cells across all 3 variants, which included only ciliated cells. A DESeq2(package v1.38.3) object was generated using the aggregated ciliated cell count matrix and COVID-19 severity score + variant group as design conditions. Log normalization of the DESeq2 count matrix was performed prior to correlation analysis.

### Statistical Testing and Data Visualization

All statistical tests were implemented in R (v4.2.3). Comparisons between cell type proportions by disease group were tested using a two-sided Kruskal-Wallis test with FDR correction across all cell types, implemented in R using the dunn.test package (v1.3.5). Spearman correlation was used where appropriate, implemented using the cor.test function in R. All boxplots were generated using the geom_boxplot() function in the ggplot2 package (v3.4.2) with the following statistics represented: center line, median; box limits, upper and lower quartiles; whiskers, 1.5x the inter-quartile range. P values, n, and all summary statistics are provided either in the results section, figure legends, figure panels, or supplementary tables. R packages dplyr (v1.1.1), tidyverse (v2.0.0), forcats (v1.0.0), stats (v4.2.3) and Matrix.utils (v0.9.7) were used for data wrangling. ggplot2 (v3.4.2), ggraph (v2.1.0), cowplot v1.1.1), ggdendro (v0.1.23), ggalluvial (v0.12.5), stringr (v1.5.0), ggrepel (v0.9.3), ggnewscale (v0.4.9), RColor brewer (v1.1-3), ggbiplot (v0.55), ggpubr v(0.6.0), gridExtra (v2.3), pheatmap (v1.0.12), ggplotify (v0.1.1), and EnhancedVolcano (v1.16.0) were used for data visualization. Color palettes were generated using iWantHue (https://medialab.github.io/iwanthue/).

## Data Availability

The annotated data can be explored via the Broad Institute Single Cell Portal (Accession: <u>SCP2593</u>) and will be available for download upon publication. Supplementary tables are available upon request.

## Code Availability

Code was written using R v4.2.3 and publicly available packages. No new algorithms or functions were created and code used in-built functions in listed packages available on CRAN. All code generated and used to analyze the data reported in this paper will be available on GitHub in the jo-m-lab repository.

## Supplementary Figures

**Supplementary Figure 1:**
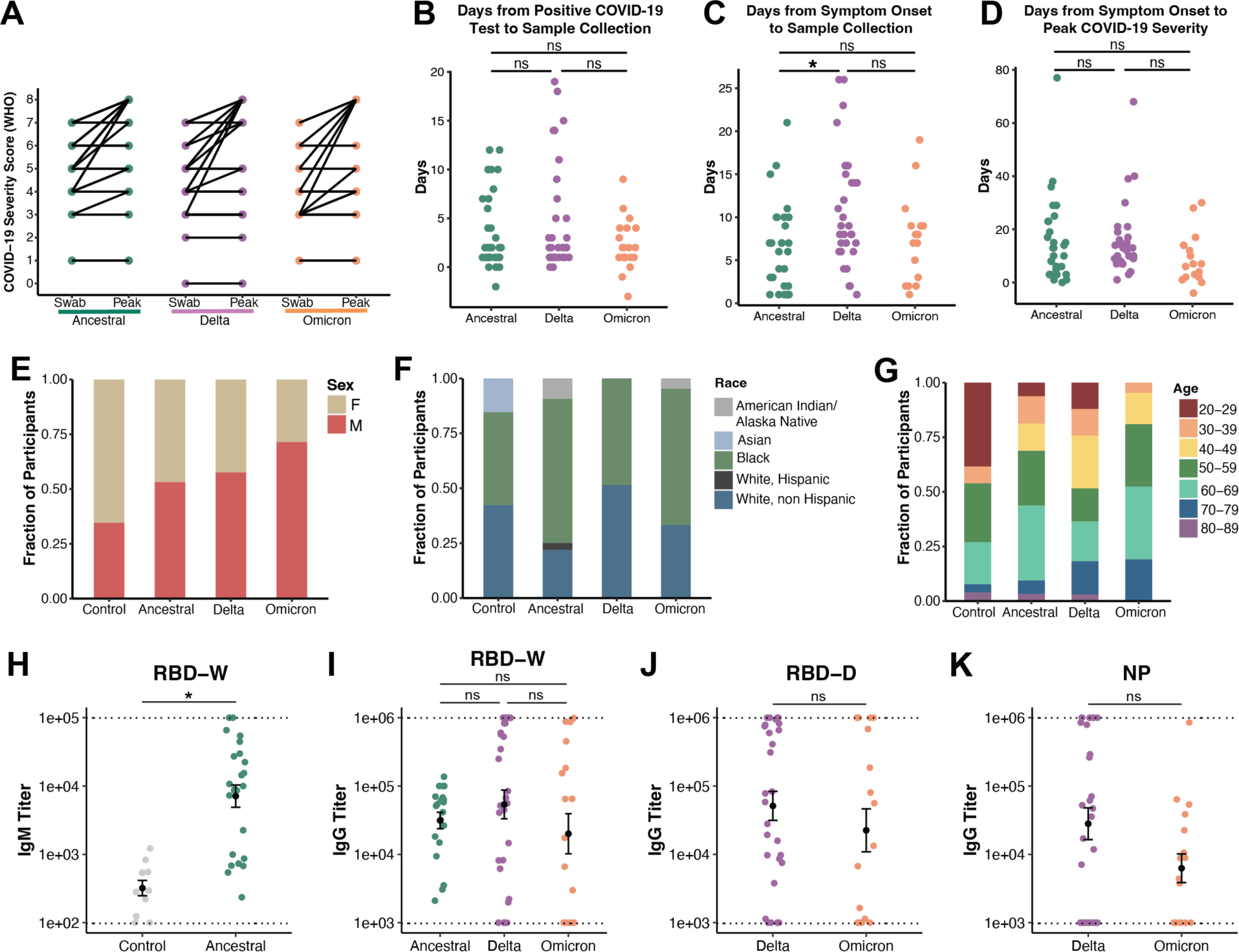
Clinical metadata by SARS-CoV-2 variant group. (A) Comparison of WHO respiratory support score at time of swab (left) and peak disease severity (right) across variant groups. (B-C) Timing of sample swab collection relative to day of positive COVID-19 test and (B) and reported symptom onset (C) (D) Duration of disease: days from symptom onset to date of highest COVID-19 severity score (WHO). Statistical test for (B-D) represents Kruskal-Wallis test results across all conditions with Benjamini-Hochberg correction for multiple comparison across all cell types. Statistical significance asterisks represent results from Dunn’s post hoc testing. *p < 0.05 (E) Participant sex by variant group (F) Participant race/ethnicity by variant group (G) Participant age by variant group (H-K) SARS-CoV-2 plasma serology across conditions. Plasma samples were taken on the same day as the nasopharyngeal swab used for scRNA-seq. Dotted lines indicate limits of detection. IgM (H) or IgG (I-K) titers against indicated SARS-CoV-2 protein separated by condition. RBD-W: original Washington strain receptor binding domain; RBD-D: Delta strain receptor binding domain; NP: nucleoprotein. Statistical test represents Kruskal-Wallis test results across all conditions with Benjamini-Hochberg correction for multiple comparison across all cell types. Statistical significance asterisks represent results from Dunn’s post hoc testing. *p < 0.05

**Supplementary Figure 2:**
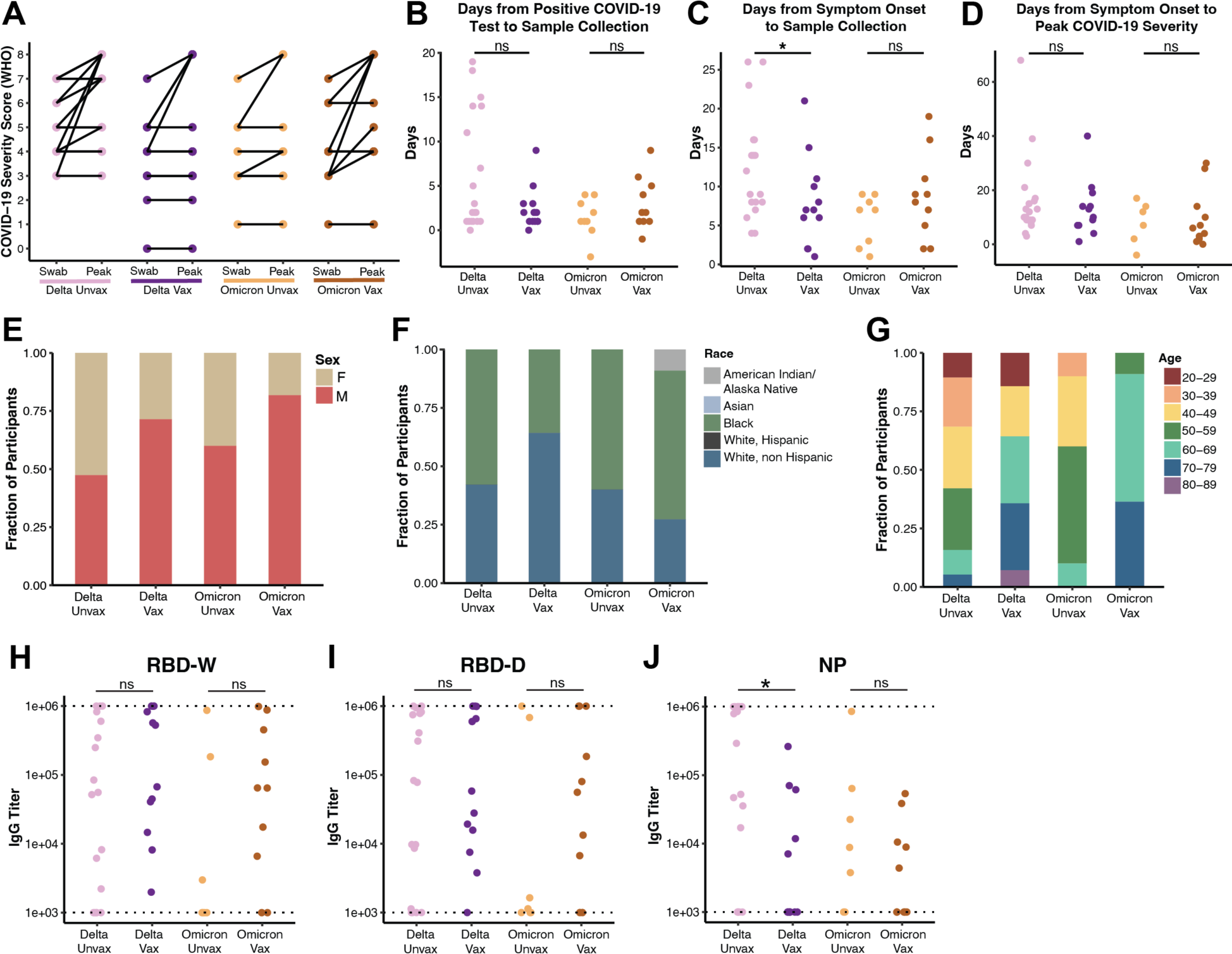
Clinical metadata by vaccination group. (A) Comparison of WHO respiratory support score at time of swab (left) and peak disease severity (right) across vaccination groups (B-C) Timing of sample swab collection relative to day of positive COVID-19 test and (B) and reported symptom onset (C) (D) Duration of disease: days from symptom onset to date of highest COVID-19 severity score (WHO). Statistical test for (B-D) represents Kruskal-Wallis test results across all conditions with Benjamini-Hochberg correction for multiple comparison across all cell types. Statistical significance asterisks represent results from Dunn’s post hoc testing. *p < 0.05 (E) Participant sex by vaccination group (F) Participant race/ethnicity by vaccination group (G) Participant age by vaccination group (H-J) SARS-CoV-2 plasma serology across vaccination groups. Plasma samples were taken on the same day as the nasopharyngeal swab used for scRNA-seq. Dotted lines indicate limits of detection. IgG titers against indicated SARS-CoV-2 protein separated by condition. RBD-W: original Washington strain receptor binding domain; RBD-D: Delta strain receptor binding domain; NP: nucleoprotein. Statistical test represents Kruskal-Wallis test results across all conditions with Benjamini-Hochberg correction for multiple comparison across all cell types. Statistical significance asterisks represent results from Dunn’s post hoc testing. *p < 0.05

**Supplementary Figure 3:**
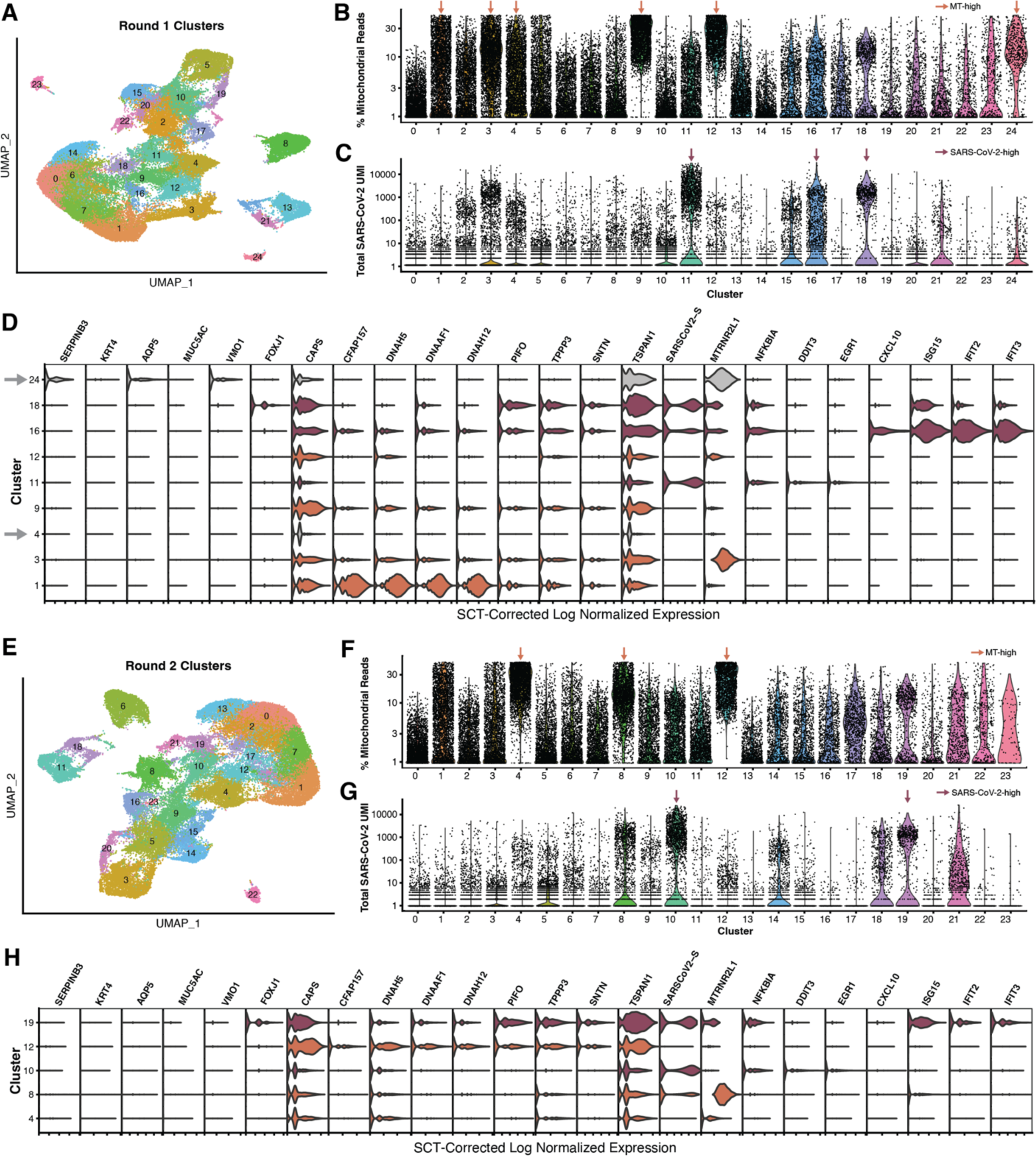
Iterative Removal of Low-Quality Cells. (A) UMAP of 55,319 cells from all participants colored by seurat cluster (B-C) Violin plots of percent mitochondrial reads (B) and total SARS-CoV-2 reads (C) for each cell, divided by clusters identified in (A). Arrows indicate clusters that were further evaluated for expression of major cell type marker genes. (D) Violin plots of cell type marker genes used to evaluate clusters identified in B-C. Gray arrows indicate clusters that were removed due to low complexity and lack of expression of marker genes (cluster 4) or due to being suspected doublets (cluster 24). See **Supplementary Table 2** for full lists of marker genes unique to each cluster. (E) UMAP of 51,314 cells from all participants colored by seurat cluster following removal of clusters specified in (D) and re-normalization (F-G) Violin plots of percent mitochondrial reads (B) and total SARS-CoV-2 reads (C) for each cell, divided by cluster identified in (E). Arrows indicate clusters that were further evaluated for expression of major cell type marker genes. (H) Violin plots of cell type marker genes used to evaluate clusters identified in F-G. No clusters were removed at this stage.

**Supplementary Figure 4:**
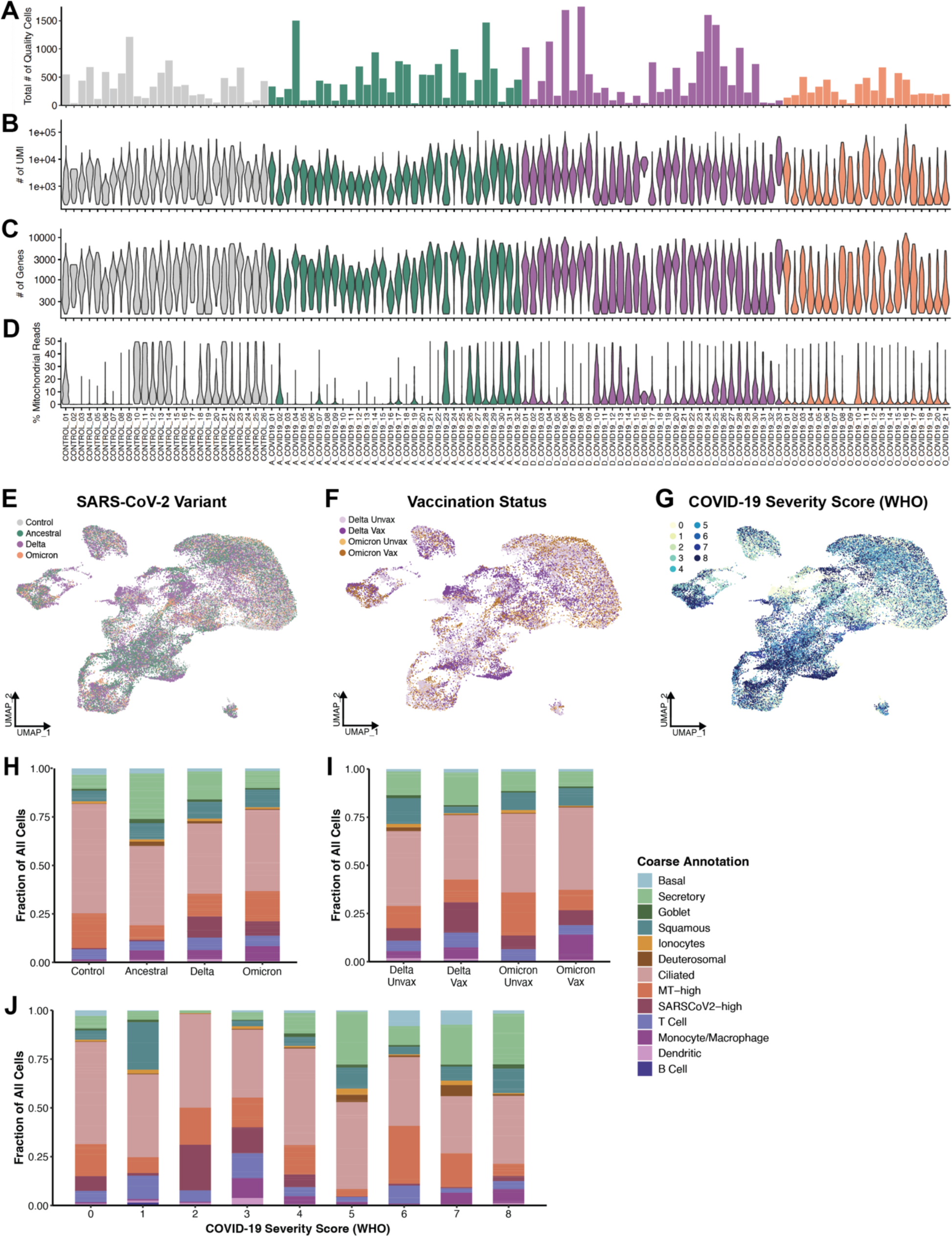
scRNA-seq Quality Metrics and Cell Type Distributions. (A) Total number of quality cells recovered for each participant (B-D) Single cell quality metrics by participant after filtering for low quality cells. UMI: unique molecular identifiers (unique transcripts) (E) UMAP of 48,730 cells from all participants colored by variant group (F) UMAP of 23, 987 cells from Delta and Omicron participants colored by vaccination status (G) UMAP of 48,730 cells cells from all participants colored by WHO peak respiratory support score (H-J) Stacked bar chart showing frequency of major cell types across variant groups (H), vaccination status (I), and COVID-19 severity (J)

**Supplementary Figure 5:**
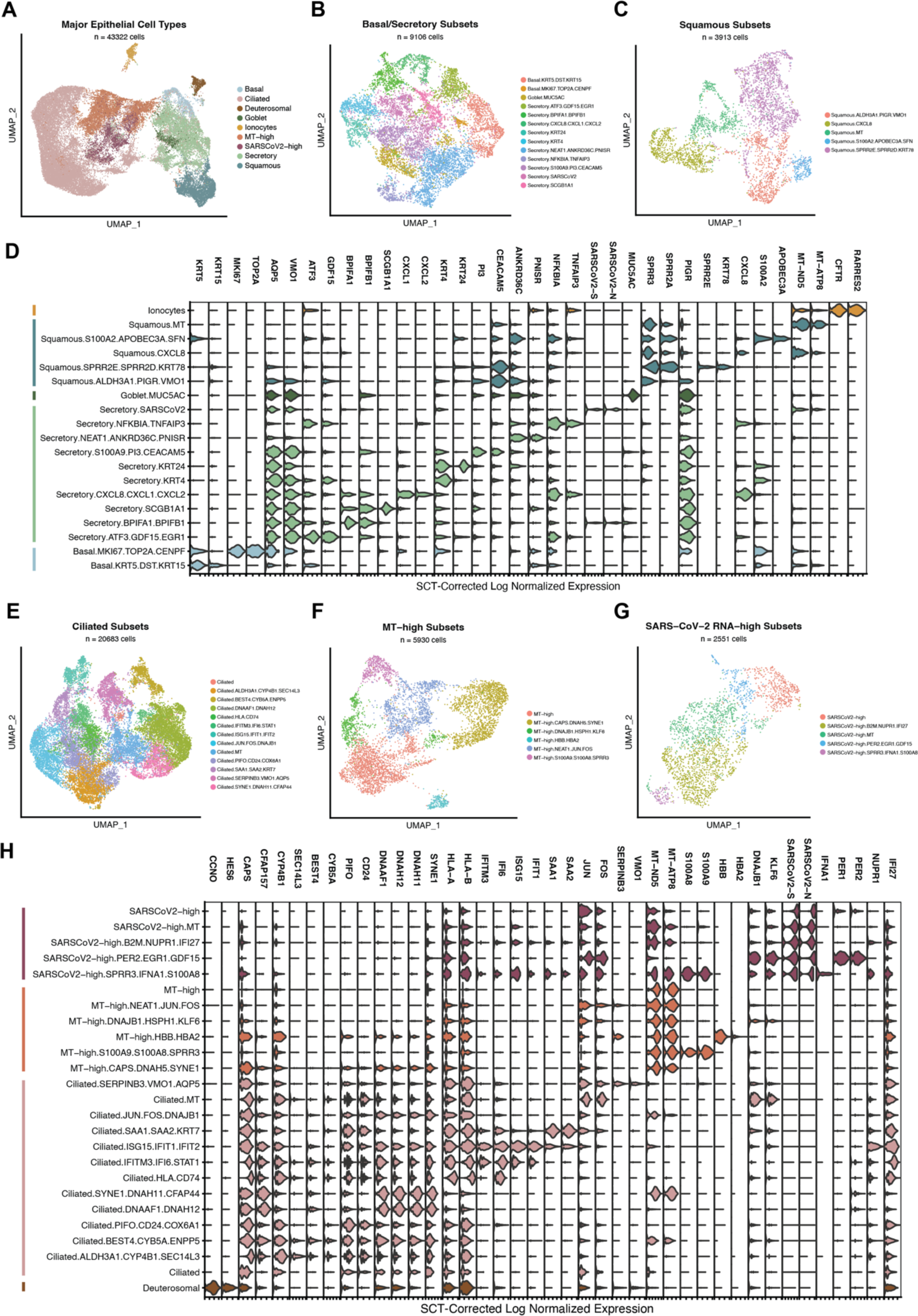
Epithelial diversity in the nasal mucosa during COVID-19. (A) UMAP of 43,322 epithelial cells from all participants colored by major epithelial cell type (defined after iterative louvain clustering) (B-C) UMAPs of detailed subclusters within basal/secretory (B) and squamous (C cells. Cells from all participants belonging to that cell type were sub-clustered with louvain clustering. See **Supplementary Tables 3-4** for full lists of cluster marker genes. (D) Violin plot of marker genes for basal, secretory, goblet, squamous, and ionocyte subsets. Colored bars represent major cell types. (E-G) UMAPs of detailed subclusters within ciliated (E), MT-high (F) and SARS-CoV-2 RNA-high (G) cells. (H) Violin plot of marker genes for ciliated, MT-high, and SARS-CoV-2 high subsets. Colored bars represent major cell types.

**Supplementary Figure 6:**
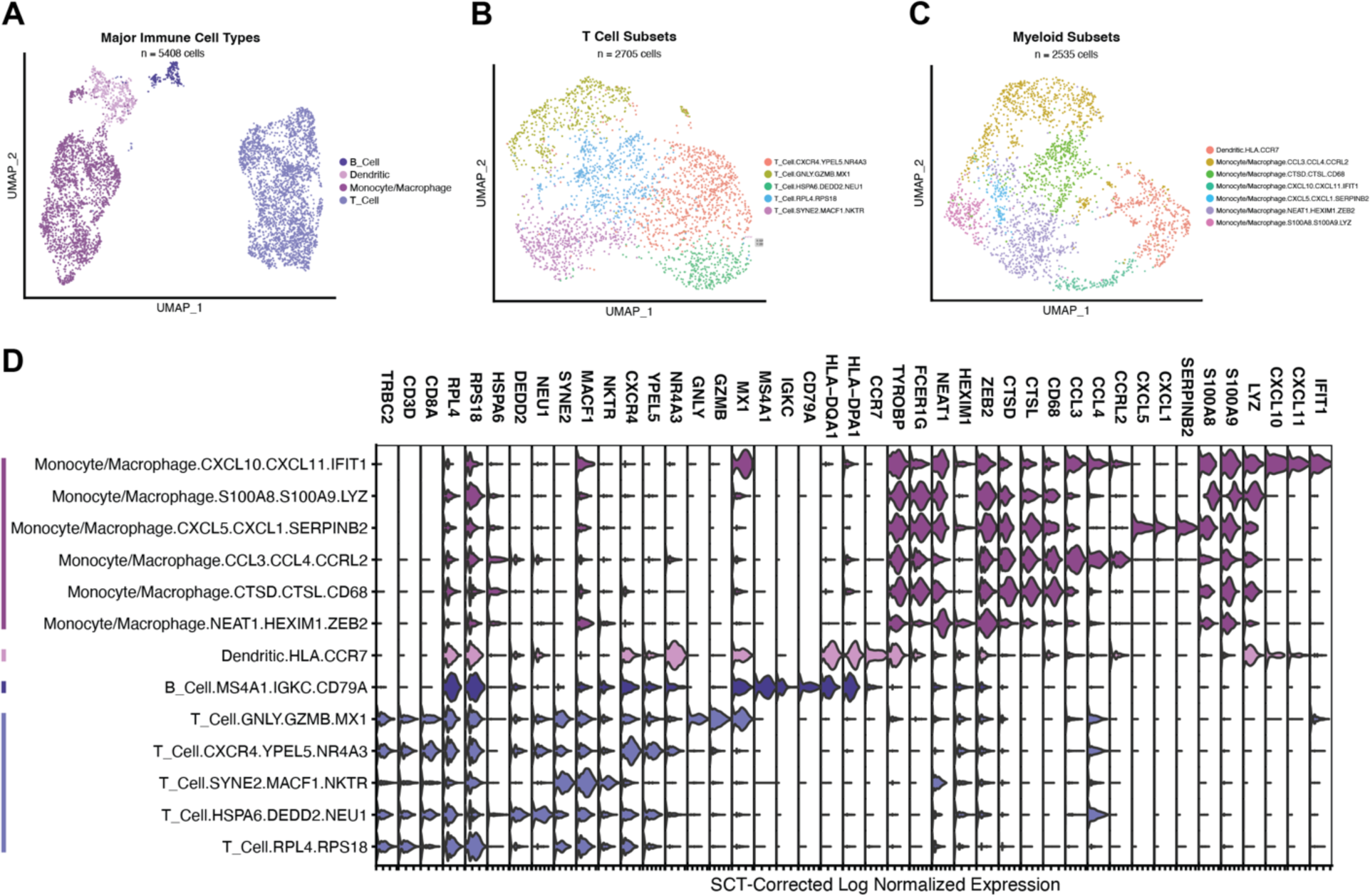
Immune cell diversity in the nasal mucosa during COVID-19. (A) UMAP of 5,408 immune cells from all participants colored by major immune cell type (B-C) UMAPs of detailed subclusters within each major immune cell type. Cells from all participants belonging to that cell type were sub-clustered with louvain clustering. (B) 2,535 myeloid cells colored by 7 detailed subsets. (C) 2,705 T cells colored by 5 detailed subsets. See **Supplementary Table 5** for full lists of marker genes. (D) Violin plot of marker genes for each detailed immune subset. Colored bars represent major cell types.

**Supplementary Figure 7:**
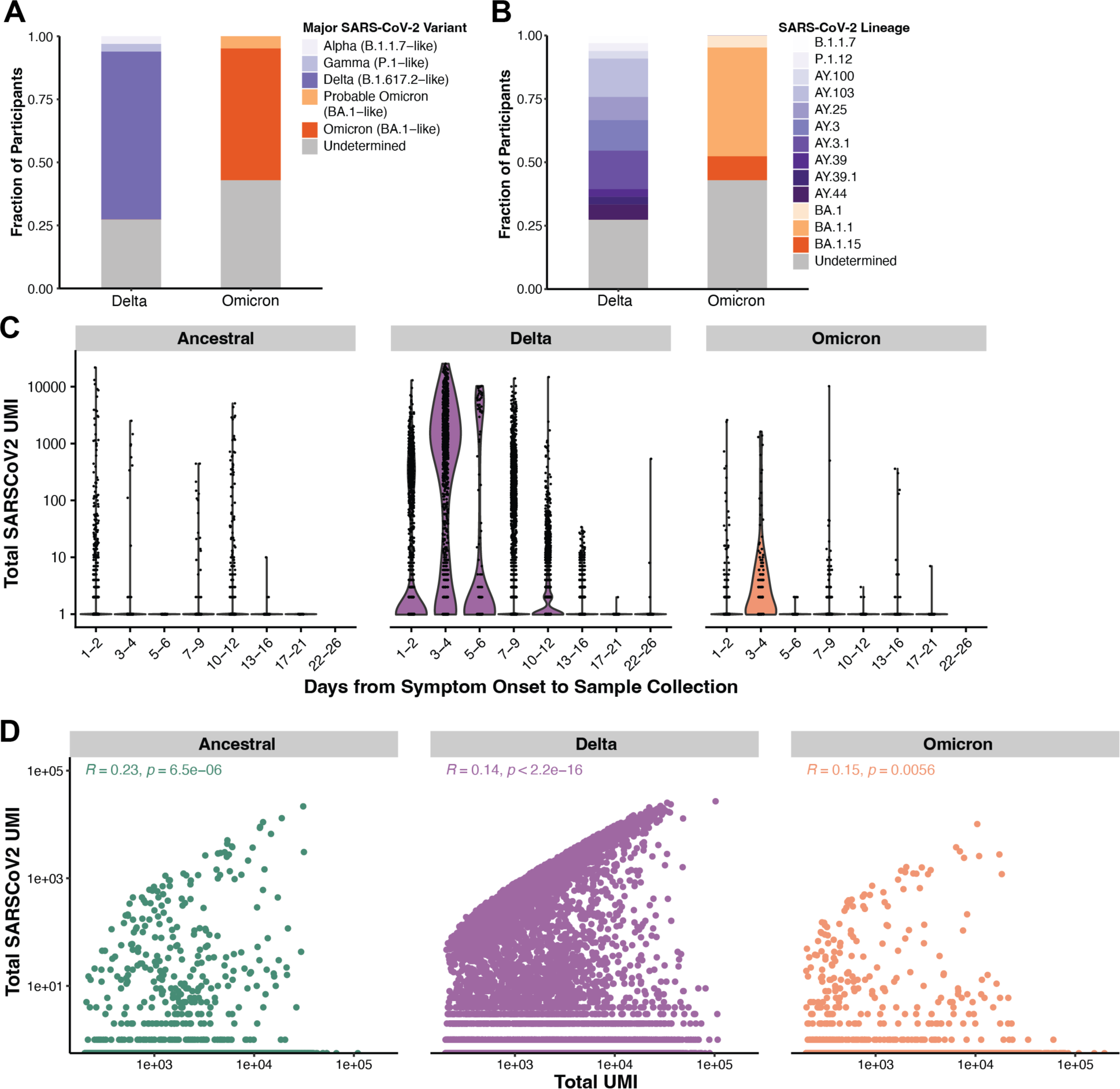
Assessment of SARS-CoV-2 RNA detection. (A-B) Results of sequencing viral RNA from supernatant of nasopharyngeal swab processing. Only samples with SARS-CoV-2 CT value <= 30 were sequenced. “Undetermined” samples are those with CT values > 30. Lineages determined using Scorpio call. (need to add more details here) (C) Violin plots of total SARS-CoV-2 transcripts detected grouped by time from symptom onset to sample collection, separated by variant. (D) Correlation of total transcripts detected in each cell with total SARS-CoV-2 transcripts detected, separated by variant. Correlations represent Spearman correlation test.

**Supplementary Figure 8:**
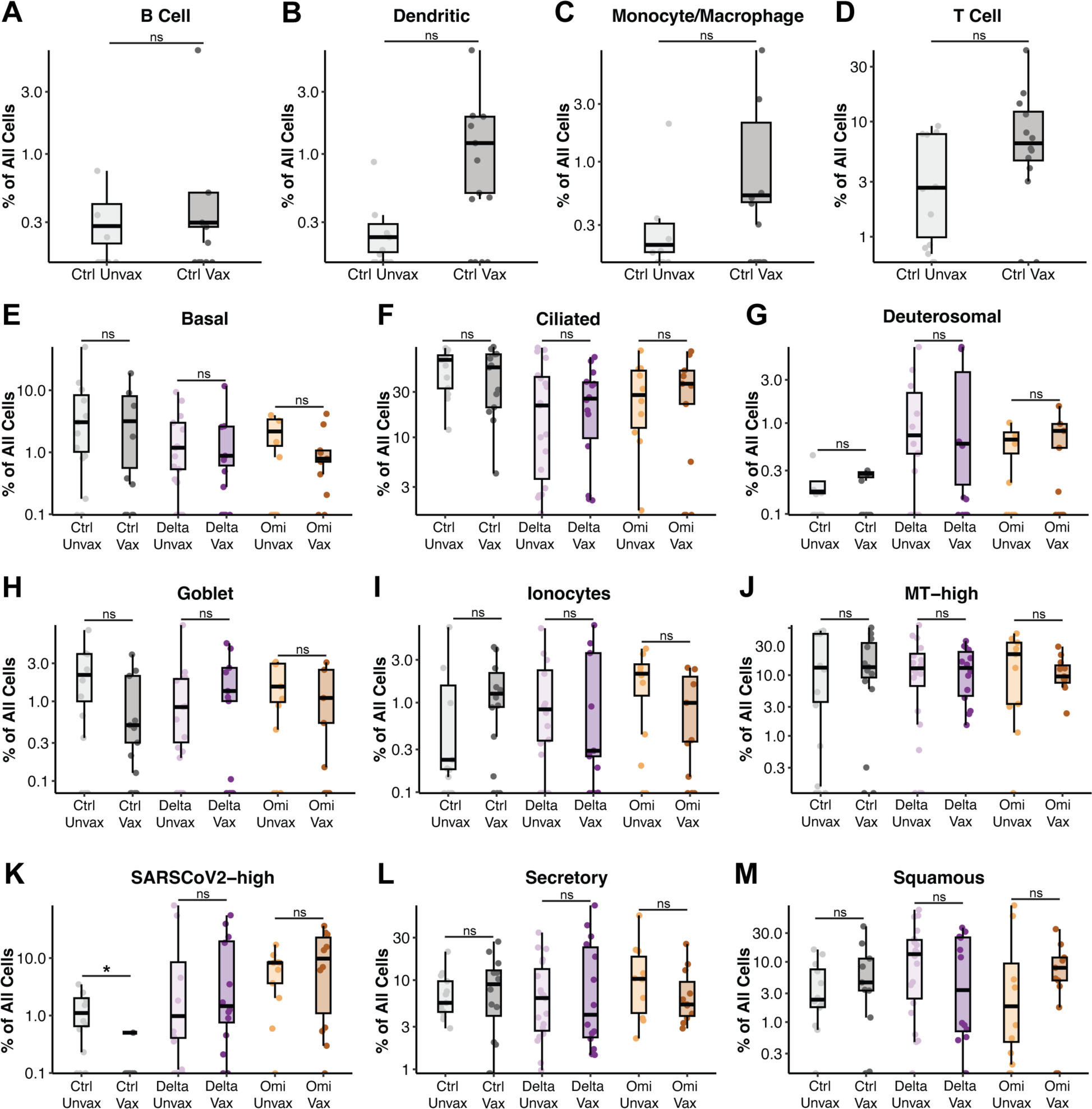
Frequency of Major Cell Types Across Vaccination Groups. (A-D) Frequency of major immune cell types as a percentage of all cells in each sample. Control unvaccinated (Ctrl Unvax) samples were collected during the Ancestral wave, and control vaccinated (Ctrl Vax) samples were collected during the Delta wave. (E-M) Frequency of major epithelial cell types as a percentage of all cells in each sample. Abbreviations: Ctrl = Control; Omi = Omicron; Unvax = unvaccinated; Vax = vaccinated. Control samples were collected as described in A-D. Statistical tests for A-M represent Kruskal-Wallis test results across all conditions with Benjamini-Hochberg correction for multiple comparison across all cell types. Statistical significance asterisks represent results from Dunn’s post hoc testing. *p < 0.05

**Supplementary Figure 9:**
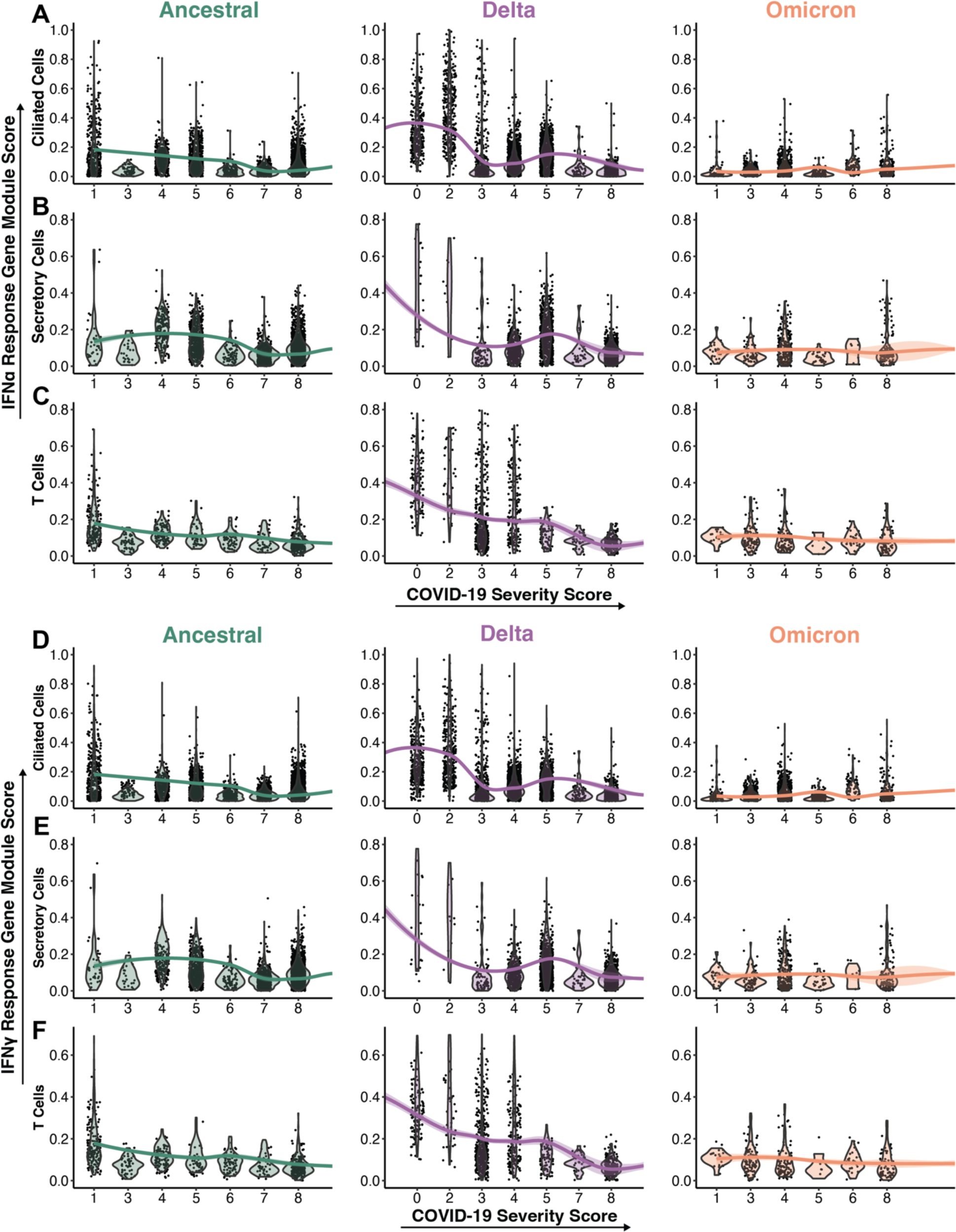
Expression of ISG Modules Across COVID-19 Severity. (A-F) Violin plots of interferon (IFN) alpha (A-C) and interferon gamma (D-F) response module score in cells from participants with varying scores of COVID-19 severity, separated by major cell type and variant cohort. Genes used for module score were derived from previously published results of stimulating human nasal basal cells with IFNa (A-C) or IFNg (D-F). See **Supplementary Table 8** for gene lists.^4^ Lines represent local regression using the locally estimated scatterplot smoothing

**Supplementary Figure 10:**
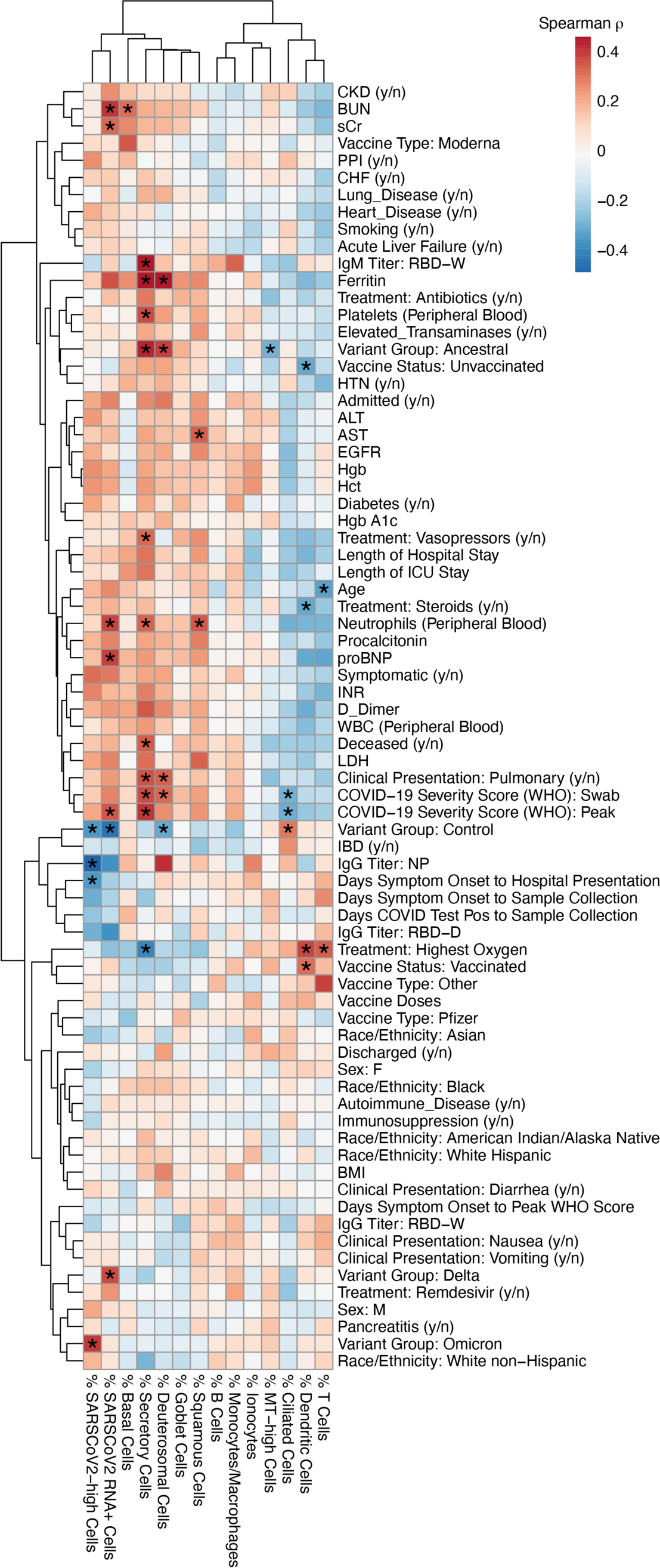
Association of clinical metadata and nasal cell type frequencies. Heatmap of spearman correlation between clinical parameters and demographic information (rows) and frequency of major cell types and SARS-CoV-2 RNA+ cells (columns). Color indicates spearman *R,* *FDR < 0.05.

## Notes

https://singlecell.broadinstitute.org/single_cell/study/SCP2593/impact-of-variants-and-vaccination-on-nasal-immunity-across-three-waves-of-sars-cov-2

